# Mechanism of translation initiation on endogenous eukaryotic circular RNAs

**DOI:** 10.1101/2025.10.31.685782

**Authors:** Philipp K. Zuber, Xueyan Li, Yifei Du, Yuliya Gordiyenko, Talal F.M. Haddad, V. Ramakrishnan

## Abstract

Eukaryotic circular RNAs (circRNAs) perform a wide variety of functions. A subset of circRNAs has been demonstrated to undergo translation *in vivo*. However, few insights exist on the underlying translation mechanisms. Here, we elucidate the basis of translation initiation in two naturally occurring circular RNAs, circMbl and circSfl. We show that *in vitro* prepared versions of both circRNAs are translated in eukaryotic cell lysates and cells. Initiation depends on the untranslated region (UTR) and can be reconstituted using only the 43S pre-initiation complex, eukaryotic initiation factors 4G, 4A, 4B, and, for circSfl, the RNA helicase DHX29. Functional assays and structural analysis of the initiation complexes suggest that scanning until recognition of the start codon follows initial ribosome landing occurring on AU-rich, accessible UTR patches upstream of the initiation sites. Together, these results provide key insights into how eukaryotic ribosomes engage with and initiate translation on circular endogenous transcripts.

## Introduction

Eukaryotic circular RNAs (circRNAs) are an abundant class of covalently closed, single stranded RNAs.^1^ Most circRNAs originate from a back-splicing reaction within a pre-mRNA, during which a 5’ splice site is connected to an upstream 3’ splice site. Usually, any introns within circRNAs are eventually spliced out, resulting in one or several circularized exon(s).^2,3^ While a fraction of back-splicing may be considered erroneous and the resulting circRNAs non-functional^4^, there is evidence for several functions being fulfilled by circRNAs. Usually, circRNAs are non-coding, and e.g. act as decoys for microRNAs^5^ or RNA-binding proteins (RBPs)^6^; regulate gene expression^6,7^; or, collectively, inhibit the dsRNA sensor PKR in absence of viral infection.^8^ Interestingly, a small subset of circRNAs is proposed to be coding and serves as templates for translation into micro-proteins.^9^ While this may not represent the prime function of most endogenous circRNAs^3^, there are well documented examples of translated circRNAs in various organisms, tissues and cell types, including human heart^10^, cancer cells^11^ mouse sperm cells^12^, and *Drosophila*.^9–13^

Among the first translatable circRNAs described was *Drosophila* circMbl(2), generated from the *muscleblind* (*mbl*) host mRNA (referred to as circMbl in the following).^9^ Interestingly, circMbl is highly conserved from flies to humans (called circMBNL1)^14^, and though no function of its translation product has been assigned, knock-down of this circRNA results in embryonic lethality or morphological defects in *Drosophila*, highlighting its importance in development.^15^ However, since circMbl also acts as a splicing regulator of its own gene locus^15^, disentangling non-coding functions of the RNA and potential functions of its translation product *in vivo* is challenging. Another translated circRNA is *Drosophila* circSfl(2) derived from the *sulfateless* (*sfl*) gene (hereafter referred to as just circSfl).^13^ CircSfl is expressed in long-lived insulin-signaling mutants and promotes longevity. The same longevity-enhancing effect was observed upon ectopic expression of the circSfl-encoded protein from linear mRNA, indicating that the phenotype is caused by the circRNA translation product.^13^ Furthermore, dozens of peptides derived from cancer-specific circRNAs were detected on the human leukocyte antigen (HLA) complex.^11,16^ These tumour cell-specific neo-epitopes allow recognition and eventual destruction of cancer cells by the immune system, and may thus provide avenues for the generation of anti-cancer vaccines.

By definition, coding circRNAs require distinct mechanisms for translation initiation compared to (linear) mRNAs. Under normal circumstances, mRNAs recruit the cap-binding eukaryotic initiation factor (eIF) 4F complex to their 5’ end. This complex consists of the scaffold protein eIF4G, eIF4E which recognizes the mRNAs’ 5’ cap, and the RNA helicase eIF4A. This “activated” mRNA can then recruit the 43S pre-initiation complex (PIC), i.e. the small 40S ribosomal subunit bound by eIFs 1, 1A, 3, 5, and a ternary complex (TC) of charged initiator tRNA (Met-tRNA^iMet^), eIF2 and GTP. Upon binding, the mRNA just downstream of eIF4F is inserted into the ribosome’s mRNA channel and the resulting 48S complex “scans” the 5’ untranslated region (UTR) to localize the start codon (annotated translation initiation site, TIS; summarized in ^19^). Scanning is promoted by the helicase activity of eIF4A, enhanced by its cofactor eIF4B^19^, or, on highly structured 5’ UTRs, by other RNA helicases such as DHX29.^20^ It was traditionally assumed that circRNAs are not translatable in principle because of their lack of a free 5’ end/cap.^21^ However, when placed in an intergenic region or circRNA context, virus-derived internal ribosome entry sites (IRESes) can mediate efficient initiation.^22,23^ This indicates that mechanistically, RNA loading into the ribosome does not necessarily require a free end. While viral IRESes are mechanistically well characterized^24^, much less is known about how endogenous coding circRNAs are activated, recruit the ribosome and localize the start codon.

Here, we present insights into the translation initiation mechanism of circMbl and circSfl. We show that *in vitro* prepared circRNA constructs can be translated in cell lysates and, at low levels, in cells. Reconstitution experiments suggest that canonical initiation factors allow recruitment of the 43S-PIC to AU-rich, accessible regions of the circRNAs UTRs. Subsequent scanning then allows identification of the TIS. These findings establish a mechanistic framework for how eukaryotic ribosomes initiate translation on natural circular RNAs, shedding light on how ribosomes access circular transcripts.

## Results

### *In vitro* generated circRNAs can be translated in lysates and in cells

To study the mechanisms of circRNA translation, we used our previously developed Trans-ribozyme based circularization (TRIC) method^23^ to generate circRNAs for an *in vitro*-based approach (Figure S1A). We chose circMbl^9^ and circSfl^13^ as model systems, since both circRNAs exhibit a size suitable for efficient characterization and have been shown to be translated *in vivo*. Both circRNAs contain the 3’ portion of the 5’ UTR and the beginning of the coding sequence (CDS) of their host mRNAs (Figure 1A). Translation was shown to initiate at the identical TIS in both mRNAs and circRNAs and terminates shortly after the ribosome has passed the back-splice junction (BSJ).^9,13^ From here onwards, we designate the annotated translated region of the circRNA its CDS and everything else as the UTR.

**Figure 1.**
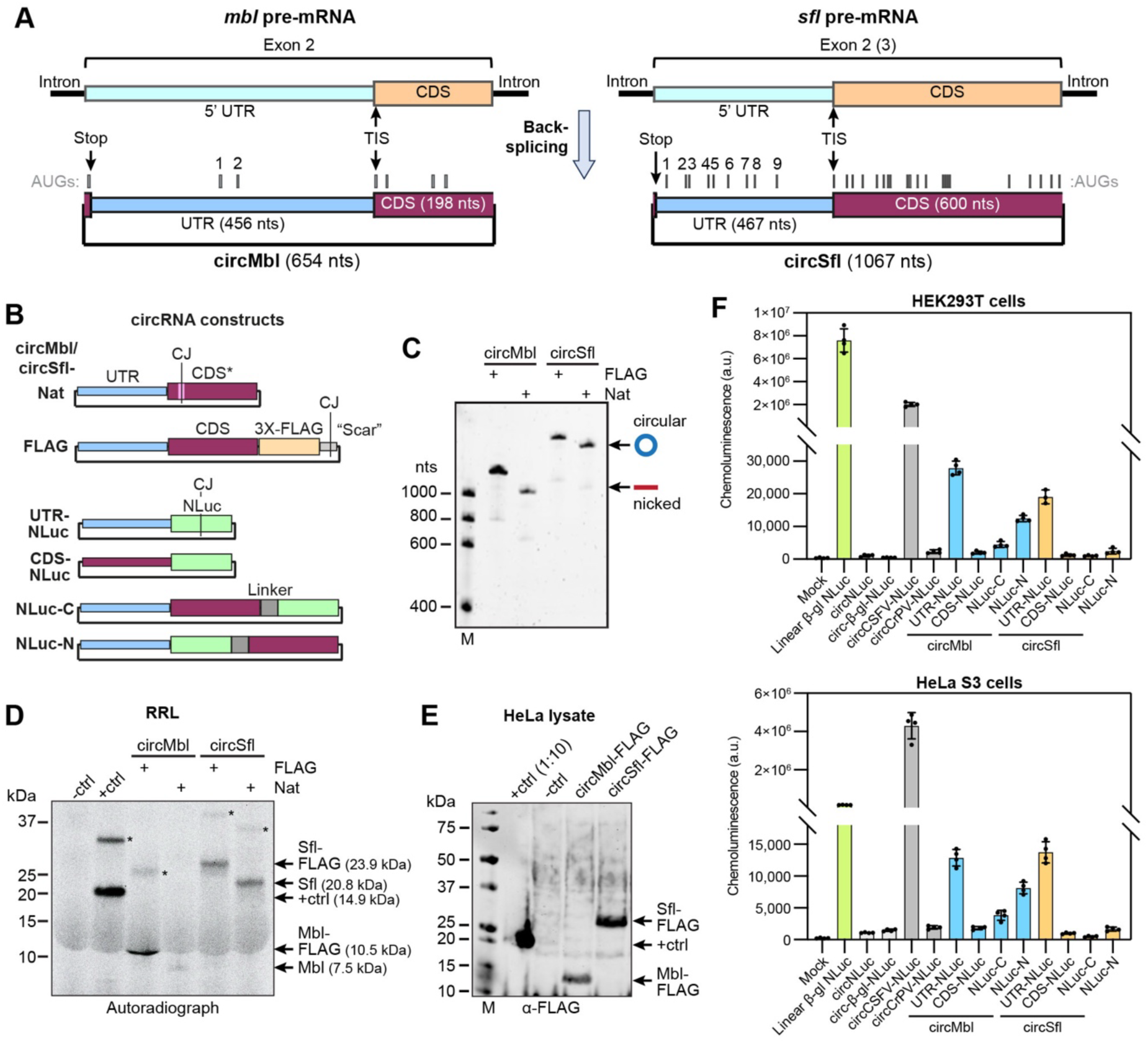
Synthetic circMbl and circSfl molecules are translated *in vitro* and in cells. (**A**) Schematic organization of circMbl/circSfl and relation to their host pre-mRNAs. Position of AUGs within the circRNA UTRs and TIS/Stop codon of the annotated CDS are indicated. (**B**) Scheme of the circRNA constructs employed in the study. The position of the circularization junction (CJ) is indicated. CDS* refers to a circRNA CDS that contains synonymous codon changes to allow *in vitro* circularization. (**C**) Urea agarose gel of circMbl/Sfl FLAG and Nat constructs. Bands corresponding to circular and nicked (linear) forms are indicated. (**D, E**) *In vitro* translation experiments of circMbl/Sfl constructs in RRL, detected by autoradiography (D), or HeLa lysate, detected by anti-FLAG Western blotting (E). A linear NLuc mRNA was used as positive control (+ctrl). The bands highlighted with asterisks are likely gel artifacts. (**F**) NLuc activity assays of HEK293T (top) or HeLa S3 (bottom) cells transfected with NLuc RNA constructs (*n* = 4).

We first prepared circRNAs closely mimicking the endogenous molecules (“Nat” constructs) and variants in which the circRNA CDS is fused to a 3X-FLAG sequence at its C-terminus (Figure 1B, C). In vitro translation in rabbit reticulocyte lysate (RRL) containing ^35^S-labelled methionine for detection revealed only one major translation product for each construct (Figure 1D), which matches the size of the peptides originating from the previously annotated CDSes. To verify that translation is not restricted to RRL, we translated the circRNA FLAG constructs in HeLa lysate, followed by anti-FLAG Western blotting (Figure 1E). In each case, one band, running close to the predicted size of the CDS-FLAG translation products was detected. We therefore conclude that circMbl and circSfl can be translated in cell lysates with translation commencing at the previously annotated TIS. However, we note that very small peptides, resulting from translation of other ORFs, may be missed by these gel-based approaches.

To quantify the translation efficiency of the circRNAs in cells, we transfected HEK293T or HeLa S3 with the circRNA constructs incorporating a nano luciferase (NLuc) CDS (Figure 1B, S1B) and quantified NLuc activity after 24 h (Figure 1F). As 5’ UTR derived sequences may more easily accommodate translation enhancing elements than CDS derived regions, we first tested circMbl/Sfl constructs containing only the corresponding UTRs along with the NLuc ORF (UTR-NLuc). Both circRNAs translated about 10-20x more than constructs containing the Cricket Paralysis Virus (CrPV) IRES, but ∼100-250x less than mediated by the Classical Swine Fever virus (CSFV) IRES or ∼60-300x less than a linear NLuc mRNA with b-globin 5’ UTR. This clearly demonstrates that both circRNA UTRs contain sequences that allow internal ribosome entry and circRNA translation in cells. Consistent with prior observations *in vivo*^9,13^, the translation efficiency is considerably lower than the levels provided by a strong viral IRES or a well translated linear mRNA. In contrast, circMbl/Sfl CDS-NLuc constructs showed a strongly reduced translational output, suggesting that the circRNA CDSes alone do not promote translation. We then tested translation of constructs that contained both the circMbl/Sfl UTR and CDS, the latter being fused to either the N- or C-terminus of NLuc. Interestingly, we observed a drastic drop in NLuc activity in the NLuc-C arrangement compared to the UTR-NLuc constructs. This disparity in apparent translational yield could e.g. be caused by differences in translation (initiation) efficiency or stability of the fusion proteins compared to isolated NLuc. Consistent with the first hypothesis, changing the position of the NLuc tag to the circRNA CDS N-terminus moderately increased the NLuc activity, indicating translation is more efficient if allowed to start at the NLuc TIS. Second, co-translational separation of NLuc from the circRNA translation product via a P2A sequence causes an increase in activity to about 50 % of the circRNA UTR-NLuc constructs (Figure S1C), showing that, the circRNA translation products indeed have a pronounced negative effect on either NLuc stability or activity when fused directly.

Quantitative reverse transcription (qRT) PCR analysis of selected transfected RNAs showed an intracellular level in HEK293T cells that was within the same order of magnitude between the different circRNAs (usually 2-3x difference) (Figure S1D). However, there was a slight inverse correlation between the observed RNA level and the length of the corresponding circRNAs that might provide an additional explanation for the differences in NLuc activity.

Together, these results suggest that (i), circMbl and circSfl show robust translation in human cells at a level comparable to that of weak viral IRESes, and (ii) the circRNA UTRs play an essential role for translation initiation. Notably, we obtained similar results when transfecting *Drosophila* S2 cells with the NLuc constructs (Figure S1E), suggesting that the generally low translation levels observed in human cell lines are not due to species-related differences in the translational apparatus.

### Circular molecules are the main source of translation

*In vitro* prepared circRNAs always contain a small portion of nicked circRNAs, i.e. linear species that result from spontaneous hydrolysis of the RNA circle (Figure 1C). We reasoned that, if occurring at a suitable position (e.g. upstream of the TIS), nicking could generate potentially translatable linear RNAs. This is especially relevant for RRL which is known for its relaxed cap-dependence^25^. To assess the contribution of nicked RNAs to the translational output, we first tested the stability of circMbl/circSfl-FLAG in RRL reactions during the *in vitro* translation experiment (Figure S2A). This showed that both circRNAs are relatively stable over the duration of a typical *in vitro* translation reaction, and that no nicked RNAs build up. Next, we deliberately nicked circMbl/Sfl-FLAG by heating under alkaline buffer conditions, resulting in different ratios of circular vs. nicked RNA (Figure S2B, D). When used for an *in vitro* translation reaction in RRL, we observed a similar level of translation at an approximate equal ratio of nicked vs. circular circMbl-FLAG compared to the starting condition (Figure S2C). This indicates that, indeed, some circMbl nicking products can be translated. However, if all initial translation is due to nicked instead of circular RNAs being used as templates, this should result in a pronounced increase in translation when the nicked:circular RNA ratio is increased, in turn suggesting that most protein synthesis occurs on circular molecules under conditions of low nicking. For circSfl-FLAG, the translational output was proportional to the amount of circRNA in the sample (Figure S2D, E). Thus, practically all translation results from the circRNA.

Overall, we conclude that one should indeed exercise caution when using *in vitro* prepared circRNAs for translation studies, especially when translational output is low, since translation of nicked RNAs is possible. However, under conditions where samples contain predominantly circular molecules, the contribution of nicked RNAs to the overall level of translation is negligible.

### CircMbl and circSfl do not bind the ribosome directly but require the complete 43S-PIC for translation initiation

Previous studies have proposed various mechanisms for translation initiation on circRNAs, such as IRES-like elements^9,17^. IRESes derived from RNA viruses employ diverse modes of internal ribosome recruitment, ranging from direct interaction with the 40S or 80S ribosome (e.g CrPV IRES) to requiring most canonical eIFs and additional IRES-trans acting factors to recruit the 43S-PIC (e.g. type-I IRESes)^24^. To assess whether the circRNAs can directly recruit the ribosome, we used a co-sedimentation assay to probe interactions with the human 40S or 80S ribosomes (Figure 2A). RNA extracted from the pellet (containing ribosomal subunits, Figure 2B) and supernatant (containing most of the unbound RNAs, Figure 2C-G) was analyzed by urea agarose gel electrophoresis. Both circMbl- and circSfl-FLAG predominantly remained in the supernatant when incubated with 40S or 80S, similar to the VHP-β negative control mRNA (Figure 2C, D). This suggests that both circRNAs do not strongly interact with isolated 40S or the 80S, and thus likely require additional factors for translation initiation.

**Figure 2.**
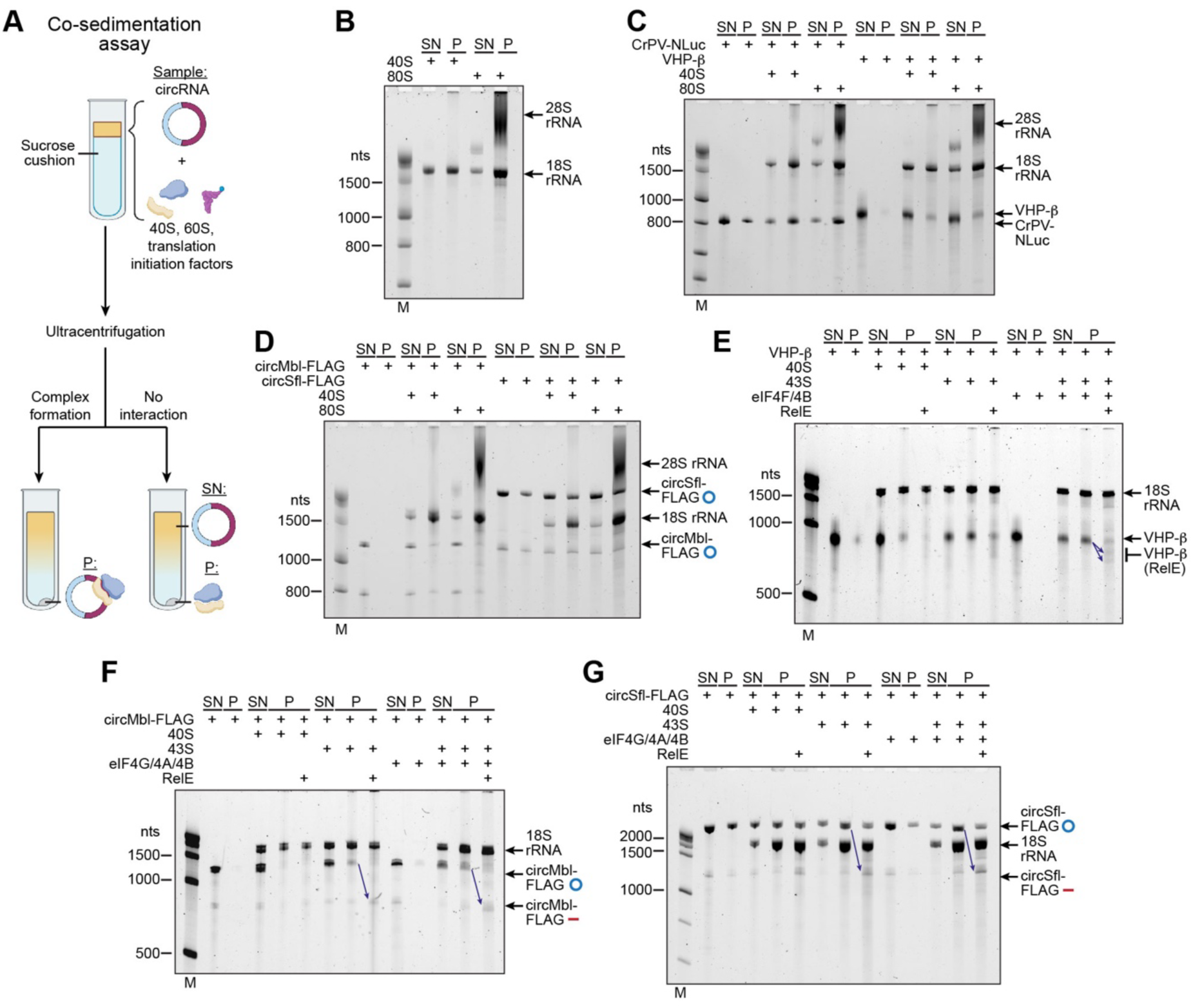
CircMbl and circSfl require the 43S-PIC for initiation complex formation. (**A**) Scheme of the co-sedimentation assay used to probe direct interaction between circRNAs and ribosomal complexes. (**B-G**) Co-sedimentation experiments of 40S/80S ribosomes or 43S complexes with the indicated RNAs, in presence or absence of eIF4F/4B or eIF4G/4A/4B. RNAs extracted from the supernatant (SN) or pellet (P) were analyzed by urea agarose gels. In (E-G), the pellet sample was incubated together with RelE to verify loading of the corresponding RNA into the ribosome (cleavage products/linearized circRNAs are indicated).

As circMbl and circSfl can be translated in RRL, we next used a RelE assay to assess initiation complex formation in the lysate. Briefly, circRNAs were incubated with RRL in absence or presence of translation inhibitors and then treated with the bacterial toxin RelE, that specifically cleaves (m)RNAs accommodated in the ribosomal mRNA channel in the A-site. RNAs are subsequently extracted, reverse transcribed using a radiolabeled primer, and cDNAs finally analyzed on sequencing gels to identify RelE cleavage sites. Negative controls (no RRL or high Mg2+ to inhibit all initiation) were used to identify background/RRL (but not ribosome) induced signal. In the presence of the non-hydrolyzable GTP analog GMP-PCP, we observed an enrichment of ribosomal complexes at the TIS and a second non-AUG site a few nucleotides upstream for circMbl (Figure 3A). For circSfl, ribosomes were mainly observed at the TIS and the next downstream AUG codon (CDS AUG-2), and to a lower extent at CDS AUG-3 (Figure 3B). In both cases, the amount and distribution of ribosomal complexes observed was independent of the construct used and is broadly consistent with the presence of one predominant *in vitro* translation product resulting from the CDS (Figure 1C). The fact that prevention of GTP hydrolysis increases the number of ribosomal complexes on both RNAs, suggests that translation initiation likely follows a pathway that requires AUG recognition by the TC, and thus most likely involves the 43S-PIC. Additionally, the presence of the nonhydrolyzable ATP analog AMP-PCP abolished RelE signals caused by the translating ribosome (marked * in Figure 3A, B), and reduced the number of ribosomes at the TIS/other AUGs to background levels, respectively, indicating that ATP hydrolysis is required for efficient ribosome recruitment.

**Figure 3.**
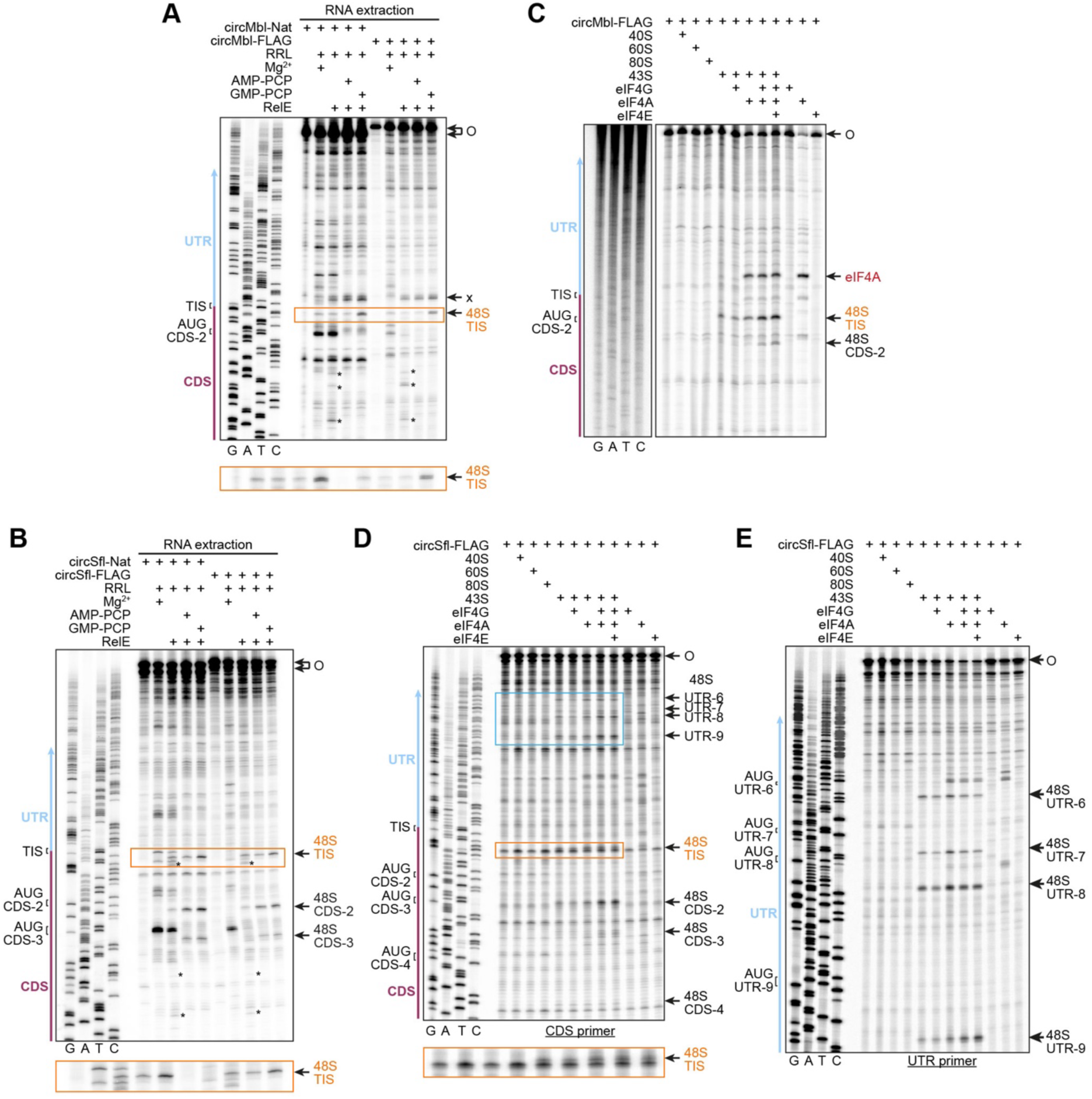
Translation initiation on circMbl/Sfl is enhanced by eIF4F factors. (**A, B**) Initiation complex formation on circMbl/Sfl-Nat and FLAG constructs in RRL, as probed by RelE cleavage assays. CircRNAs were incubated in absence or presence of RRL (+/- inhibitors), treated with RelE, RNAs extracted, and cleavage positions visualized by primer extension assays. The x in (A) marks a RelE cut at a non-AUG position upstream of circMbl TIS. Asterisks mark RelE cuts by the translating ribosome. (**C-E**) Initiation complex formation on circMbl-FLAG (C) and circSfl-FLAG (D, E) in a reconstituted human system, assayed by toe-printing. For circSfl, primers binding to the CDS (D) or UTR region (E) were used. Complex formation minimally requires the 43S-PIC and is enhanced by eIF4F or eIF4G/4A.

### The eIF4G/4A complex increases translation initiation on circMbl and circSfl

To further identify putative factors required for translation initiation on circMbl and circSfl in addition to the 43S-PIC, we purified 43S/48S complexes from GMP-PNP-supplemented RRL followed by mass spectrometry analysis. This revealed the presence of 43S-PIC factors (i.e. 40S proteins, eIF1/1A/2/3/5), the eIF4F complex (eIF4G/4A and low amounts of eIF4E), and various RBPs and RNA helicases (Table S1). To assess the contribution of the identified factors to initiation on circMbl and circSfl, we employed ribosome toe-printing assays based on a reconstituted human translation initiation system. Here, the factors to be tested are incubated with RNA, and ribosomal complexes are then detected on sequencing gels as read-through stops of reverse transcription, initiated from a radiolabeled primer annealed downstream of the site of interest. Consistent with the co-sedimentation experiments, we did not observe any extra bands upon addition of the ribosome subunits (Figure 3C-E). However, when adding the 43S-PIC, we obtained a weak toe-print at the TIS for circMbl-FLAG (Figure 3C), and toe-prints at the TIS, CDS AUG-2 (Figure 3C) and the four AUGs upstream the TIS (UTR AUG 6-9, compare Figure 1A) for circSfl-FLAG (Figure 3E).

We next tested whether eIF4G and/or eIF4A, or the eIF4F complex would improve initiation complex formation. Indeed, addition of eIF4G/4A markedly stimulated complex formation at the circMbl TIS (Figure 3C) and the CDS AUG-2 and UTR AUGs 6-9 of circSfl (Figure 3D, E). Interestingly, the increase in 48S formation is caused even by intact eIF4F, showing that the presence of the cap-binding protein eIF4E does not interfere with the role of eIF4G/A even though no 5’ cap is present here.

To test whether the initiation enhancing effect of eIF4F factors is due to an increase in recruitment efficiency or post-recruitment activities, such as promoting scanning of the circRNA for the start codon, we again employed a co-sedimentation assay. Additionally, we treated the pelleted ribosomes with RelE to check whether RNAs were loaded into the ribosome’s mRNA binding channel. For VHP-β mRNA, this assay revealed clear co-sedimentation with the 43S-PIC in the presence of eIF4F/4B, and two RelE induced cuts (Figure 2E), showing the functionality of the assay (notably, under the test conditions, there was also a significant co-sedimentation with the 43S-PIC alone). Consistent with the toe-printing assays, we observed a moderate but significant increase in complex formation for circMbl- and circSfl-FLAG when the 43S-PIC was added, compared to 40S alone. Furthermore, the extent of co-sedimentation was indeed further enhanced when eIF4G/4A and eIF4B (see below) were added (Figure 2F, G). This indicates that eIF4F factors at least partially function through enhancing recruitment of the ribosome to the circRNAs.

Importantly, since both circMbl and circSfl were linearized when RelE was added to the sedimented complexes, the observed initiation complexes indeed form on the intact circRNAs. Additional evidence indicates that the same is true under toe-printing conditions: (i) circMbl-/Sfl-FLAG remained stable under toe-printing conditions (Figure S3A). (ii) Usage of an RT with reduced RNAse H activity caused the “full-length” signal (i.e. reverse transcription around the RNA circle for one round) to disappear and a signal in the well, generated by rolling circle reverse transcription, to appear (Figure S3B). (iii) Deliberate nicking of the circRNAs decreased the amount of initiation complexes formed (Figure S3C-F).

### eIF4G/4A and eIF4B cooperate to promote ribosomal scanning on circMbl

We next asked whether any additional factors, such as the RBPs or RNA helicases identified circRNA initiation complexes purified from RRL (Table S1), would be able to further improve circMbl/Sfl translation efficiency and start codon selection *in vitro*. Therefore, we prepared recombinant eIF4B (a co-factor of eIF4A^26^), DHX29 (RNA helicase required for 48S scanning through highly structured 5’ UTRs^20^), cytoplasmic polyA binding protein (PABP), YBOX1 (RBP with multiple functions, e.g. regulation of translation initiation^27^) and PA2G4 (RBP implicated in IRES-mediated translation. ^28^) for further testing. When added to circMbl-FLAG along with eIF4A/4G, we observed an increase in 43S-PIC recruitment in the presence of eIF4B; all other factors were agnostic to translation initiation (Figure 4A).

**Figure 4.**
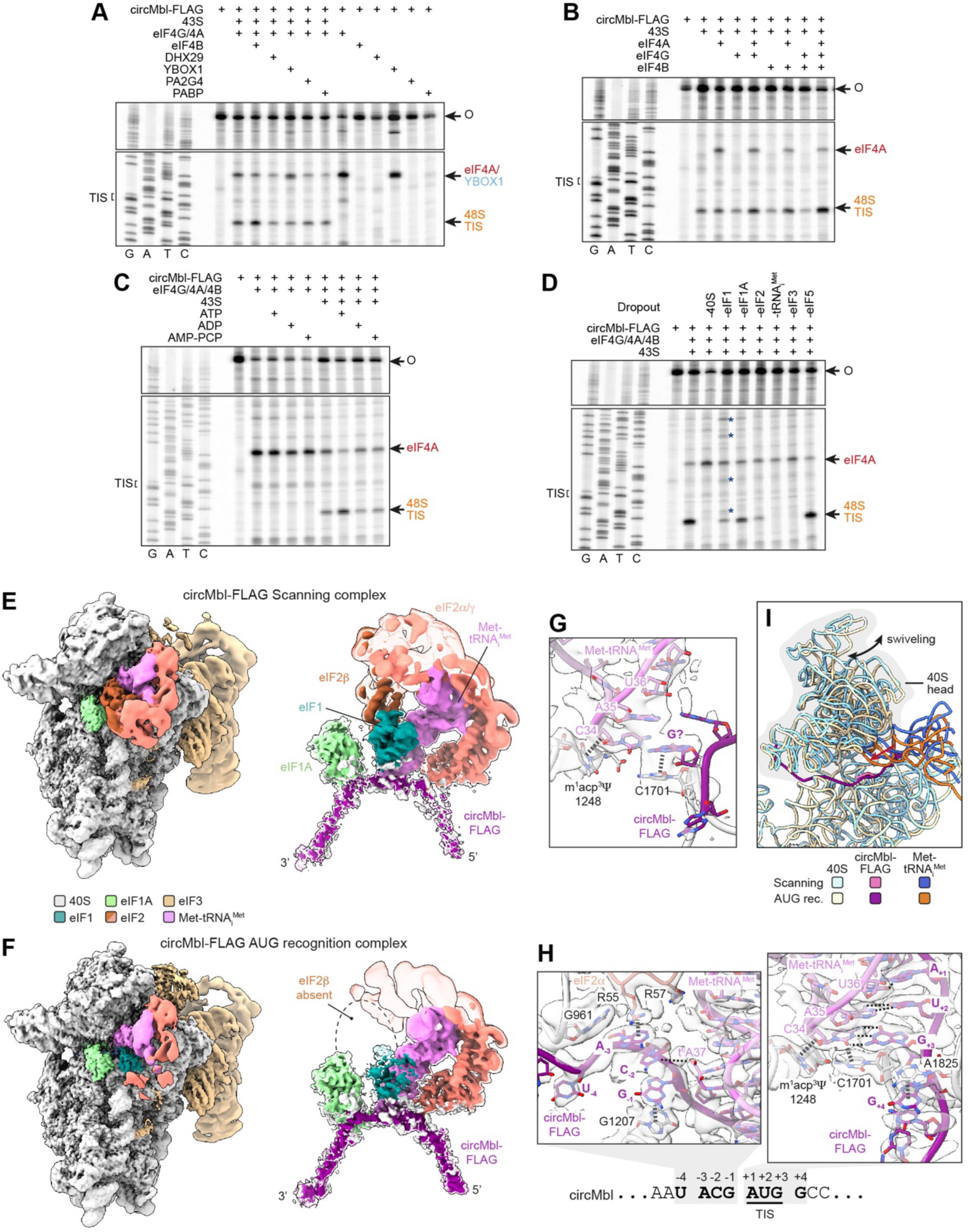
Translation initiation on circMbl involves ribosomal scanning. (**A-D**) Ribosomal toe-printing assays probing 48S complex formation on circMbl-FLAG in a reconstituted system. Toe-prints of 48S complexes and of individual factors are indicated. Initiation complex formation is increased by eIF4B (A), which acts in synergy with eIF4G and eIF4A (B). 48S complex formation is dependent on ATP hydrolysis (C) and requires all 43S-PIC factors except for eIF5 (D). (**E, F**) CryoEM maps of the circMbl-FLAG 48S complex in the scanning (E) or AUG recognition states (F), depicting the overall composition (left) or the TC, eIF1/1A and RNA region (right; map is shown at high [non-transparent] and low thresholds [transparent]). Components are colored as indicated. (**G, H**) Close-up of the tRNA_i_^Met^:circRNA interaction in the scanning (G) or AUG recognition complexes (H). Components are colored as in (E, F). (**I**) Superposition of the scanning and AUG recognition complexes, highlighting differences in orientation of the 40S head (18S rRNA), Met-tRNA_i_^Met^ and the circRNA. Components are colored as indicated.

To further investigate the interplay between eIF4G, 4A and 4B, we repeated the toe-printing assay on circMbl-FLAG using all possible combinations of the three factors (Figure 4B). In isolation, only eIF4A caused a small increase in initiation complexes formed at the TIS. Combining eIF4A with either eIF4G or eIF4B led to a further increase in initiation, while combining all three factors together resulted in a further marked increase. This is consistent with the ability of both eIF4G and eIF4B to stimulate the ATPase and RNA helicase activity of eIF4A^26^. Additionally, ATP hydrolysis was required for the stimulatory effect of eIF4A, since initiation was only increased by ATP but markedly reduced in its absence, or by ADP or AMP-PCP (Figure 4C).

Finally, we were interested whether initiation complex formation on circMbl-FLAG required all, or only a subset of factors that constitute the 43S-PIC. Dropout experiments revealed that omitting the 40S, initiator tRNA and eIF3 completely abolished, while absence of eIF1, 1A and 2 (strongly) reduced 48S complex formation at the TIS (Figure 4D). Only dropout of eIF5 had no influence on initiation. Interestingly, 43S-PICs lacking eIF1 caused a trace of new toe-prints upstream of the TIS (asterisks in Figure 4D), indicating the recognition of non/near cognate start codons and consistent with eIF1’s function to increase stringency of start codon selection.^19^

Together, these results support a model in which the 48S complex scans the circRNA UTR region in an eIF4A/ATP-dependent manner to locate the start codon.

### Cryo-EM captures scanning and AUG recognition states of circMbl initiation complexes

To gain deeper mechanistic insights into the translation initiation on circMbl, we reconstituted its translation initiation complex *in vitro* using circMbl-FLAG, eIF4G/4A/4B and the 43S-PIC, and determined its structure by cryo-EM (Figure S4, Table S2). Classification revealed two major ribosomal states, (i) a scanning complex (Figure 4E; S5A), and (ii) the ribosome in the AUG recognition state (Figure 4F; S5B). In both complexes, we identified most of the components, except for eIF4G/4A (Figure S5C, D), eIF4B and eIF5.

The scanning complex shows clear density for eIF1, eIF1A, Met-tRNA_i_^Met^, and the three eIF2 subunits (Figure 4E) - hallmarks of a 43S-PIC that has not engaged in start codon recognition. The tRNA_i_^Met^ is positioned between the P_IN_ and P_OUT_ states, as has been observed before (Figure S5E ^29^), and only loosely interacts with RNA in the mRNA channel (Figure 4G). The circRNA density in the RNA binding channel is poorly defined, consistent with the absence of specific interactions with the circRNA and suggesting multiple positions of the ribosome along the circRNA. The 40S head in the scanning complex adopts a swiveled conformation (Figure 4I, S5F), similar as described previously.^29^

The AUG recognition complex shows reduced density for eIF1 and eIF2γ, with eIF2β absent (Figure 4F). This is caused by conformational changes following start codon recognition, which eventually cause eviction of eIF2 from the complex, along with eIF1.^30^ Unlike in the scanning complex, there is complete pairing of the tRNA_i_^Met^ anticodon and an AUG codon (Figure 4H), and the 40S head is in a fully closed conformation with the tRNA_i_^Met^ in the P_IN_ state (Figure S5F). Due to stabilizing interactions with 18S rRNA residues, tRNA_i_^Met^ and eIF2α, the four nucleotides upstream and one nucleotide downstream of the AUG codon are well resolved (Figure 4H). This allows unambiguous assignment of the RNA sequence and suggests that most ribosomes in the AUG recognition state are stalled at the TIS, consistent with the toe-printing experiments.

Overall, the visualization of distinct scanning and AUG recognition states provides direct structural evidence that ribosomes can engage and scan circular RNAs to faithfully recognize the TIS.

### Translation initiation on circSfl requires DHX29-driven scanning

Like circMbl, translation initiation complex formation on circSfl-FLAG minimally requires the 43S-PIC and is enhanced by eIF4G/4A or eIF4F. However, in circSfl-FLAG 48S complexes localize not only to the TIS, but also the four UTR AUG codons upstream of the TIS and the second/third codon of the CDS (Figure 3D, E). In the reconstituted sample, the relative occupancy at the TIS is thus lower compared to what is observed in RRL (Figure 3B), prompting us to investigate whether additional factors could improve translation initiation and enhance fidelity of start codon selection. To this end, we performed the factor screening experiment for circSfl-FLAG (Figure 5A). This revealed that eIF4B markedly increased the amount of initiation complexes formed, however mostly at the UTR AUGs. Strikingly, the addition of DHX29 improved both the efficiency and specificity of TIS recognition. As DHX29 is known to function most efficiently on linear mRNAs when the initial loading of the 43S-PIC is mediated by eIF4B^20^, we examined the interplay between eIF4B and DHX29. Indeed, the simultaneous presence of eIF4B and DHX29 together with eIF4A/4G further improved both the absolute amount of initiation complexes forming at the TIS (Figure 5B). Together, these results suggest that translation initiation on circSfl-FLAG requires eIF4G/4A/4B-promoted loading of the 43S-PIC on the circRNA and subsequent DHX29 driven 48S scanning through RNA structures to reach the start codon.

**Figure 5.**
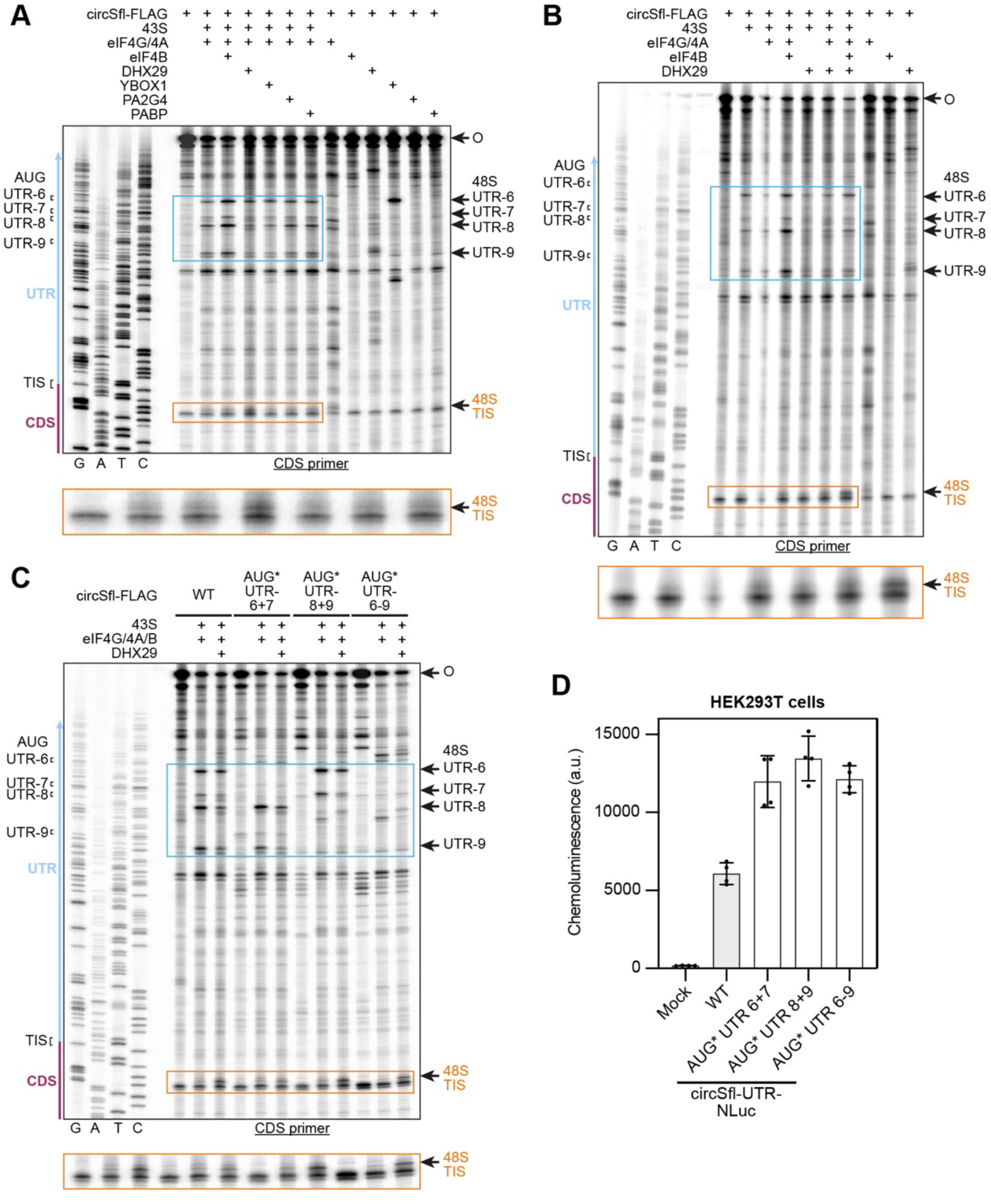
Efficient recognition of the TIS in circSfl requires DHX29. (**A-C**) Toe-printing assays on circSfl-FLAG in a reconstituted system testing influence of additional factors (A), interplay between eIF4B and DHX29 (B) and the influence of mutations of AUG UTR 6+7, 8+9 and 6-9 (C). The region around the UTR AUGs is highlighted (blue box). The area around the TIS is shown enlarged. (**D**) NLuc activity assays of circSfl-UTR-NLuc variants containing mutations of UTR AUGs 6+7, 8+9 and 6-9 in HEK293T cells (*n* = 4).

The formation of initiation complexes in the circSfl UTR is homologous to a situation where upstream ORFs in an mRNA regulate the translation of the main ORF. We thus wondered whether translation initiating at the UTR AUGs 6-9 also (negatively) regulates translation of the circSfl CDS. Therefore, we devised double mutants of the AUGs UTR 6+7, and 8+9, and constructs in which all four AUGs were mutated in either the circSfl-FLAG or circSfl-UTR-NLuc contexts. Toe-printing showed that all mutations abolished the existence of 48S complexes at the corresponding UTR AUGs (Figure 5C). Cell transfection of the circSfl-UTR-NLuc mutant constructs showed an about two-fold increase in NLuc activity for all three UTR AUG deletions Thus, these findings indicate that upstream AUGs within the circSfl UTR act as negative regulators of translation from the main CDS, likely by trapping scanning ribosomes. Moreover, they highlight a critical role for DHX29 in overcoming structural barriers and ensuring accurate and efficient translation initiation on structured circular RNA templates.

### Putative 43S-PIC landing sites localize to AU-rich regions of the circRNAs

The observation that the UTR region drives the bulk of translation initiation events on circMbl and circSfl (Figure 1F) suggests the presence of discrete RNA elements that serve as recruitment or landing sites for the 43S-PIC. To identify possible 43S landing sites, we probed circMbl/Sfl-FLAG with SARS CoV-2 Nsp1. Amongst other functions, Nsp1 cleaves host mRNAs in a 43S-PIC/eIF3g dependent manner to shut down translation. On capped mRNAs, Nsp1 cuts can be observed from the very 5’ end^31,32^, while on viral IRESes (e.g.CrPV and EMCV IRESes), the most 5’ Nsp1 cleavage events localize to the region that is first inserted into the ribosome during initiation.^32^ We thus reasoned that Nsp1 cleavage sites could serve as a proxy for potential ribosome landing sites on circRNAs. To test this idea, we assembled Nsp1-“poisoned” 43S complexes, and added circMbl/Sfl-FLAG along with eIF4G/4A/4B. After incubation, RNAs were then extracted, reverse transcribed and the cDNAs analyzed on sequencing gels (Figure 6A, B). This revealed two weak but specific cuts in the circMbl UTR (Figure 6A) and several cuts in both the circSfl UTR and CDS (Figure 6B).

**Figure 6.**
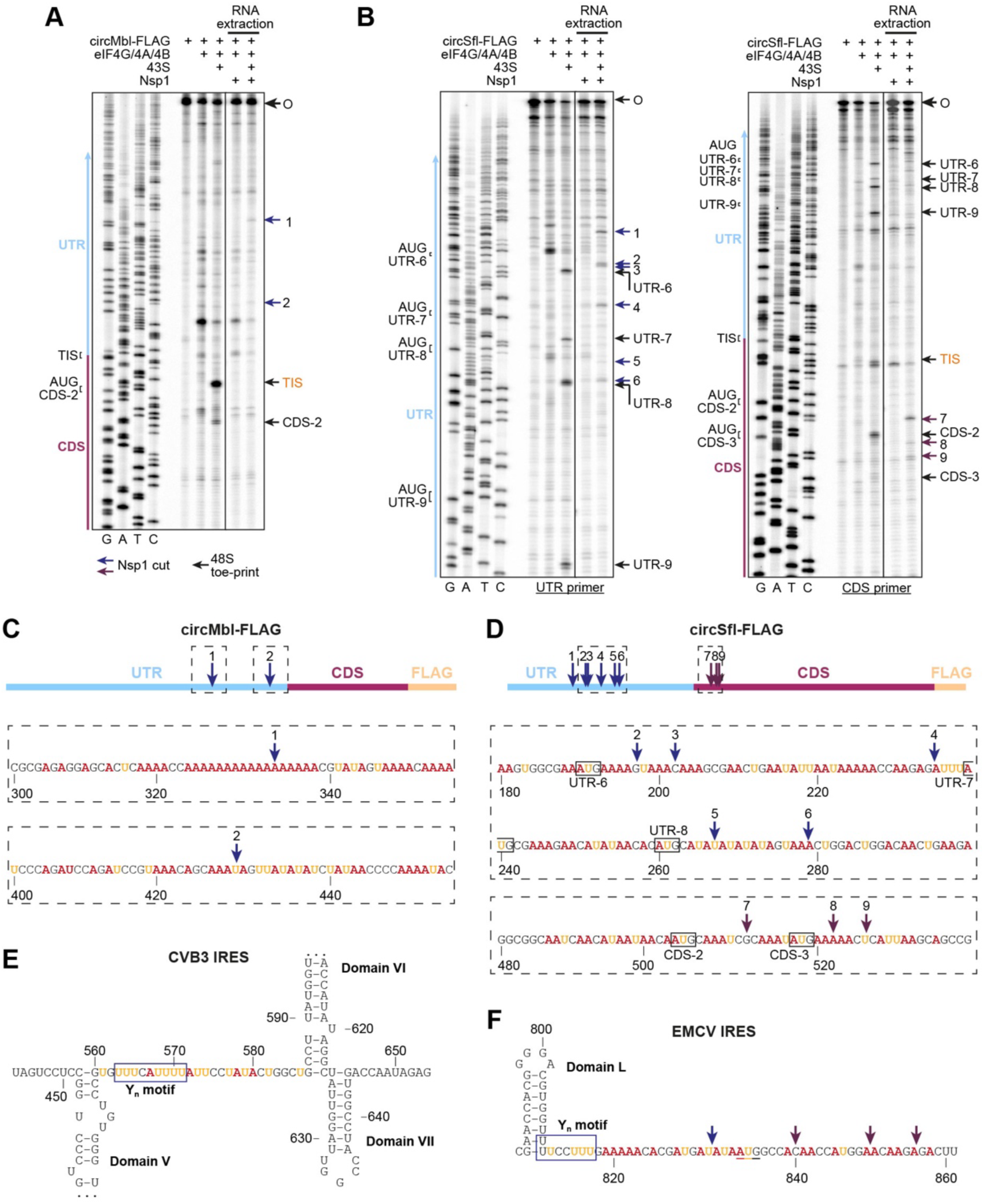
Ribosome landing on circMbl and circSfl occurs on AU-rich RNA regions. (**A, B**) Ribosomal toe-printing and Nsp1 cleavage assays probing 48S complex formation and 43S-PIC landing on circMbl-FLAG (A), and circSfl-FLAG (B) in a reconstituted system. Toe-prints of 48S complexes (black arrows) and Nsp1-induced cuts (UTR: blue arrows, CDS: magenta arrows) are indicated. For circSfl-FLAG, RT primers annealing to the UTR (B, left) or CDS (B, right) were used. (**C, D**) Position of Nsp1 cleavage sites (arrows) determined in (A, B) on circMbl-FLAG (C) or circSfl-FLAG (D). Top: Scheme of the circRNA regions, drawn to scale. Bottom: Sequence around Nsp1 cleavage sites (regions boxed above). Adenine and uracil residues are highlighted. (**E, F**) Sequence and secondary structure of the CVB3 (**E**) and EMCV (**F**) IRESes. Structural domains, the conserved pyrimidine-rich (Y_n_) motifs and adenine/uracil residues with and adjacent to it are highlighted. Nsp1 cleavage cuts from ref. ^32^ are shown as arrows in (F).

In circMbl, the two cuts localize to regions ∼50 or 140 nt upstream of the TIS (Figure 6C). Interestingly, this region also showed additional signals of 48S complexes being stalled at non-cognate codons upon omission of eIF1 (Figure 4D), indicating that the ribosome lands in the region just upstream of the TIS and subsequently scans downstream.

For circSfl-FLAG, most cuts localize to the region around UTR AUGs 6–8 (Figure 6B), which is consistent with 48S formation and downstream scanning in this region. The cuts in the circSfl CDS (Figure 6C) might be an additional explanation for the formation of initiation complexes downstream of the TIS, other than leaky scanning. We note that, while our experiments are not able to distinguish Nsp1 cuts resulting from individual landing events, or movement of the ribosome along the RNA and subsequent, delayed cleavage, the fact that signals occur at distant sites (>100 nt apart) likely indicates several independent ribosome entry sites, suggesting redundancy.

We finally wondered whether the putative ribosomal landing sites localize to any specific sequence motifs or structures within the circRNAs. Interestingly, most Nsp1 cuts are in regions enriched for adenine and uracil residues, but depleted for guanine (Figure 6C, D). Considering that AU-rich sequences are usually less likely to form stable secondary structure, these A-rich tracts might resemble the ribosome recruitment platforms in cardiovirus and enterovirus IRESes. The latter are composed of a conserved Y_n_ motif and an often AU-rich but more variable spacer that together act as flexible docking platforms for 43S-PIC recruitment (Figure 6E, F).^33^ In analogy to these viral IRESes, we therefore propose that accessible, AU-rich patches of RNA in the UTR serve as the landing sites for the 43S-PIC on the circRNAs.

Overall, these results suggest that translation initiation on both circRNAs features landing of the 43S-PIC on accessible, adenine-rich stretches of RNA, predominantly located in the UTR regions. Following landing, the 48S complex then scans downstream to localize the TIS.

## Discussion

### Translation of natural TRIC-generated circRNAs *in vitro* and in cells

Studying the function of natural, endogenous circRNA is challenging. *In vivo* and in cells, detailed investigations are complicated by low expression levels of individual circRNAs^2,3^, and usually require overexpression from potentially imperfect circularization constructs.^18^ This is especially problematic for studying circRNA translation, since linear splicing by-products could be translated and mistakenly interpreted as circRNA translation.^34^ Here, we have used our previously established TRIC method^23^ to generate circMbl and circSfl (Figure 1A), and study their translation capabilities and the underlying mechanisms of translation initiation *in vitro*.

Cell lysate-based translation assays showed that both circRNAs are translated into one major product (Figure 1D, E) and RelE assays in RRL showed initiation complex formation at the expected TIS (Figure 3A, B), thus confirming the previously annotated ORFs. Cell transfection assays (Figure 1F) reveal that circMbl or circSfl UTRs promote NLuc translation at a level similar to that enabled by the CrPV IRES, yet remained ∼100–250-fold lower than translation driven by the more potent CSFV IRES or a well-translated linear β-globin–NLuc mRNA. This places circRNA UTR-mediated initiation in a range comparable to weak viral IRESes, but well below that of canonical cap-dependent mRNAs. This relatively low translation level is consistent with *in vivo* results, such as low association of e.g. circMbl with polysomes, or weak bands in Western blots for mbl^9^ or α-FLAG-sfl.^13^ Interestingly, addition of the circRNA CDSes has an apparent negative effect on translational output of the NLuc constructs (Figure 1E). Consistent with a low stability of circRNA translation products^17^, this effect is at least partially caused by a lower stability of NLuc-mbl/sfl fusions, since trimming off the circMbl/Sfl translation product from NLuc via a P2A sequence markedly increased translation (Figure S1E). As circRNAs containing the full sequence more closely mimic the natural molecules than UTR-only constructs, this strategy is useful to accurately assess the amount of initiation events.

While our circRNAs are highly purified, even extensive treatment with exonuclease does not fully remove nicked species due to persistent, spontaneous hydrolysis of the circle (Figure 1C). This raises the concern of nicked RNAs being translated in lysates or even in cells. Indeed, when intentionally nicking circMbl-FLAG, the translation level observed at a ∼1:1 ratio (circular:nicked) in RRL was approximately comparable to that of unnicked sample (Figure S2B, C), indicating that some translation does result from nicked RNAs. However, the contribution was much less than expected if all translation observed in the initial sample was templated by nicked RNAs. For circSfl, the contribution of nicking to the overall translation was much smaller (Figure S2D, E). Together, this shows that if nicking is kept low (and the lysate is sufficiently cap-dependent), the translation observed can almost entirely be attributed to the circRNA.

### Proposed mechanism for translation initiation on circMbl and circSfl

Our interaction studies and toe-printing assays indicate that the minimal ribosomal complex able to be recruited to the circRNAs is the 43S-PIC (Figure 2F, G, 4D). The amount of initiation complexes forming at circMbl/Sfl AUG codons is further enhanced significantly by the eIF4F, or eIF4A/4G. This factor requirement is consistent with a previous eIF knock-down screen performed for circSfl^35^, which identified a strong dependence of circSfl translation on eIFs 1, 1A, 2, and 3, suggesting that initiation requires the 43S-PIC, as well as eIF4G and 4A (note that the *Drosophila* DHX29 homolog was not tested).

Interestingly, the finding that presence of eIF4E did not influence the level of initiation complex formation on both circMbl and circSfl (Figure 3C-E) suggests that even the intact eIF4F complex could engage in circRNA translation. However, recent single molecule studies show that yeast eIF4F is actively removed from cap-distal positions on mRNAs though the ATPase activity of eIF4A, but stabilized when bound at the cap, promoting “activation” of the mRNA^36^. This suggests that the eIF4F complex would eventually select against association with circRNAs, and thus actively mitigate translation of cap-less RNAs. Conversely, we speculate that conditions where association of eIF4E with the cap or eIF4G is inhibited or reduced, e.g. during certain cellular stresses, would result in a higher chance of association of the 43S with endogenous circRNAs. Consistent with this idea, down-regulating cap-dependent translation in *Drosophila* by starvation, leads to enhanced translation of circMbl.^9^

On both circMbl and circSfl, the initiation-stimulating activity of eIF4G/4A was further enhanced by eIF4B (Figure 4A, 5A). How do eIF4G/4A and eIF4B promote translation initiation on circRNAs? Our interaction studies and toe-printing assays indicate that they likely act through a combination of (i) recruiting/stabilizing the 43S-PIC on the circRNA and (ii) enhancing scanning of the resulting 48S towards the TIS. First, the amount of formed 48S complexes on the circRNAs is significantly increased by eIF4G/4A/4B (Figure 2F, G). This suggests that the eIF4F factors assist in recruiting the 43S-PIC to, and/or stabilize the complex on the circRNAs. In line with this, both eIF4A and eIF4G are potent RBPs that bind throughout the 5’ UTR and gene body of mRNAs.^36^ While not explicitly tested here, it is possible that eIF4A preferentially binds certain sequence motifs that could facilitate recruitment to specific sites in the circRNAs. Second, several lines of evidence indicate that efficient recognition of the TIS involves eIF4A-driven ribosomal scanning: (i) eIF4A drives initiation complex formation, and is stimulated by eIF4G and/or eIF4B (Figure 4B), both known to promote scanning through increasing eIF4A’s ATPase and RNA helicase activity.^26,37^ (ii) Enhancement by eIF4A/4G/4B is ATP-hydrolysis dependent (Figure 4C). (iii) Omission of eIF1 from the 43S-PIC results in a “trace” of new 48S toe-prints upstream of the TIS in circMbl-FLAG (Figure 4D), resulting from relaxed stringency of start site selection. Additionally, the fact that DHX29 is required for efficient 48S formation at the circSfl TIS (Figure 5A, B) suggests that active modulation of the circRNA secondary structure is necessary for the ribosome to access the start codon.

How does the 43S-PIC recognize suitable landing sites with the circRNAs? Using probing with Nsp1, which cleaves mRNAs and viral IRESes upon loading into the ribosome^32^, we identified several candidate sites within the circRNAs UTRs and beginning of the CDS (Figure 6A, B). Most sites display a particularly high AU-content and are largely depleted for guanine residues (Figure 6C, D). It is conceivable that these regions would show a higher degree of flexibility than more G-rich sequences and would thus constitute an accessible patch for landing of the 43S-PIC and insertion of the RNA into the ribosome. This is homologous to the Y_n_ motifs of type-I or type-II IRESes, that together with a downstream spacer provide a “flat” landing site for the 43S-PIC in-between/after structured IRES domains.^38^

Overall, we therefore propose the following mechanism for translation initiation on circMbl and circSfl (Figure 7): 1. probing of the circRNA for accessible regions by the 43S-PIC, aided by stabilizing interactions with RNA bound eIF4G/4A, and 2. subsequent, eIF4A/4G/4B or DHX29 driven scanning to localize a start codon in a suitable sequence context.

**Figure 7.**
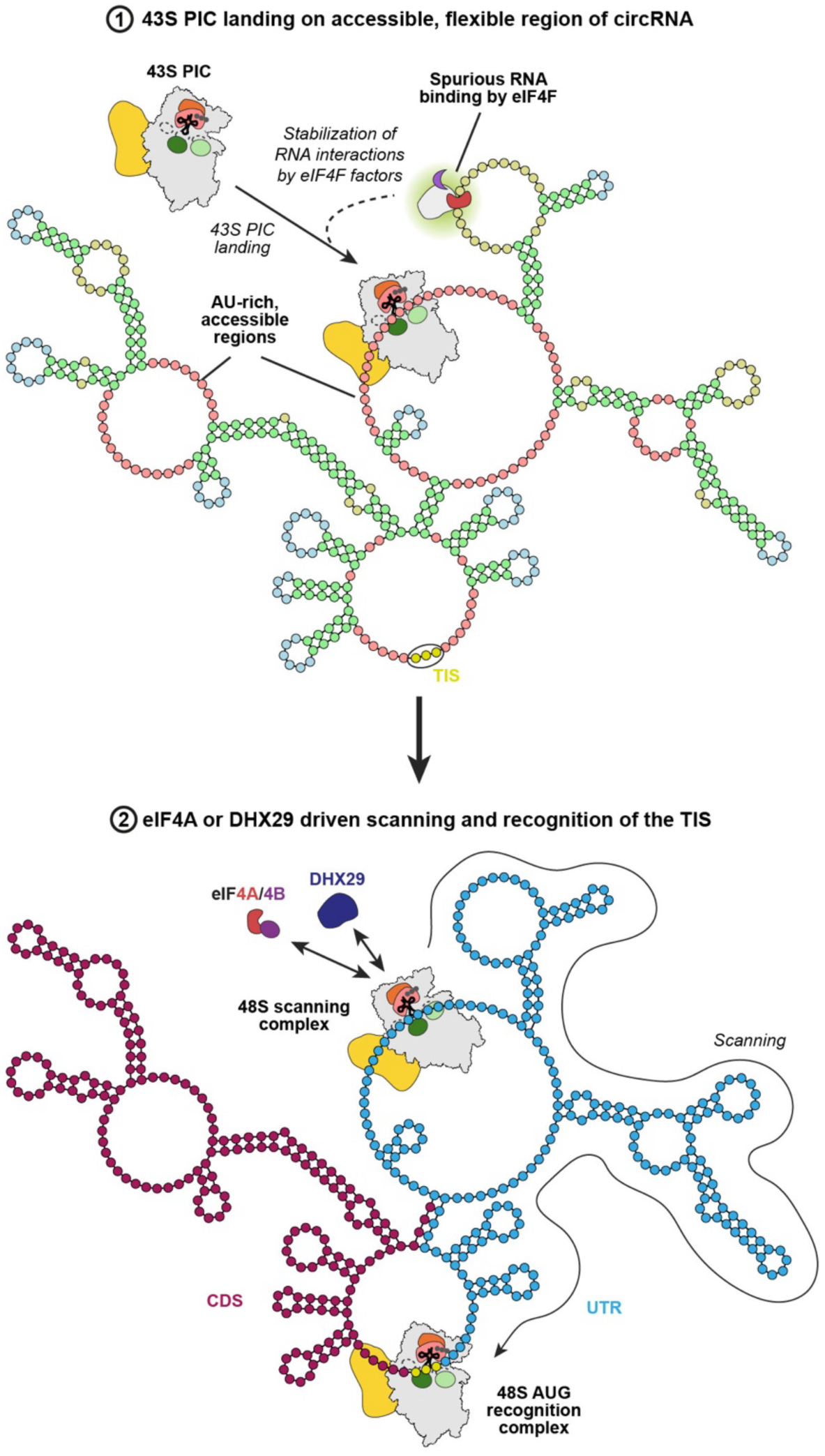
Model for translation initiation on natural circRNAs. We propose that the 43S-PIC recognizes AU-rich, accessible regions within endogenous circRNAs (red in step 1), occurring in e.g. 5’ UTR derived sections. Recruitment may be enhanced through stabilizing interactions with the eIF4F complex. The loaded 48S complex can scan the circRNA in 3’ direction through either the helicase activity of eIF4A (enhanced by eIF4G and/or eIF4B) or, in case of strong secondary structures, DHX29 (step 2) until recognition of a suitable start codon.

### Comparison of circRNA translation initiation to other modes of cap-independent translation initiation

We wondered whether the mechanism described here may be applicable to other putative translatable circRNAs. An analysis of the ribosome-associated circRNAs from *Drosophila*^9^ shows two main classes: circRNAs containing stretches of the host mRNA 5’ UTR and CDS (thus containing the mRNA’s TIS, exemplified by circMbl/circSfl), and those consisting only of CDS exons (Figure S6A). We reason that CDS regions are unlikely to contain “classical” IRESes akin to those of viruses (i.e. highly folded structures) since this would induce two-fold evolutionary pressure: (i) on the host mRNA amino acid sequence and (ii) on the IRES structure. Analysis of the translatable 5’ UTR+CDS circRNAs revealed that 5’ UTR-dervied regions span a wide range of sizes (7 – 1200 nts, median 274 nts); some obviously too small to contain virus-like IRESes (Figure S6B). While we cannot exclude that some 5’ UTR derived regions contain structured regions that contribute to translation initiation, it is conceivable that both 5’ UTR and CDS-derived regions would exhibit sufficiently accessible, flexible RNA stretches that would allow low levels of translation initiation to happen. For CDS-only circRNAs, the apparent translational yield may additionally be amplified by rolling circle translation, thus increasing the apparent translational yield.^17^

Translation initiation on coding circRNAs is conceptually similar to internal initiation provided by viral IRESes. Those are typically characterized by highly folded RNA structures, that harness non-canonical interactions with the translation initiation machinery. The employed mechanisms and interactions range from direct recruitment of the 40S/80S ribosome (e.g. CrPV IRES), to requiring all canonical eIFs and additional factors (type I IRESes).^24^ While a lot of viral IRESes directly place the start codon in the ribosomal P-site, type-I IRESes use a combination of eIF4G/4A-mediated ribosome recruitment followed by ribosome scanning to localize the TIS.^33^ However, all classical viral IRESes direct the ribosome to a precise landing site, sometimes formed by conserved pyrimidine-rich motifs.^24,38^ This well-defined and efficient ribosome recruitment contrasts with the mechanism proposed here, which may more rely on the ribosome’s intrinsic preference to localize on accessible, unstructured AU-rich regions, helped by more labile RNA association of eIF4G/4A in the vicinity. Interestingly though, some plant RNA viruses contain adenine rich, unfolded intergenic regions in their genomes, functionally akin to the unfolded elements on circMbl/Sfl. Consistently, the translational output of these viral IRES-like elements is comparable to the translation level seen for the circRNAs studied here.^39^

In conclusion, our study provides a simple mechanism for translation initiation on endogenous circRNAs, that features 43S-PIC docking onto flexible, AU-rich elements aided by eIF4F factors, followed by ribosomal scanning.

## Acknowledgments

We thank Dr Vish Chandrasekaran for helpful discussions and advice; Dr Yanyi Jiang for critical comments on the manuscript; the Electron Microscopy and Scientific Computing facilities of the MRC LMB for support and access to instruments; Dr. Keren Turton for maintaining the MRC LMB insect cell facility and for providing cells; Dr Catarina Franco and Dr Farida Begum of the MRC LMB Biological Mass Spectrometry and Proteomics Facility for mass spectrometry sample preparation and analysis. P.K.Z was supported by Postdoctoral Fellowships from the German National Academy of Sciences Leopoldina (LPDS 2021-14) and EMBO (ALTF 778-2021). X.L. was supported by an EMBO Postdoctoral Fellowship (ALTF 937-2022). Y.D. was supported by the China Postdoctoral Science Foundation (PC2021083) and the Leverhulme Trust (ECF-2022-525). VR was supported by the UK Medical Research Council (MC_U105184332) and the Wellcome Trust Senior Investigator award (WT096570).

## Author contributions

P.K.Z., X.L., Y.D. and V.R. conceived the study. P.K.Z, X.L., Y.D., Y.G., and T.H. performed the experiments. P.K.Z., Y.D. and X.L. wrote the manuscript with input from all authors. V.R. supervised the research and helped edit the manuscript.

