## Supplemental Information for "Mechanism of translation initiation on endogenous eukaryotic circular RNAs"

### Materials and Methods

#### Molecular cloning

The sequences of *D. melanogaster* circMbl(2) (dme_circ_0001328) and circSfl(2) (dme_circ_0001693) were retrieved from circBase [1]. The design of circularization constructs used in this study was based on the TRIC approach as described in [2]. Briefly, *in vitro* transcription templates for circMbl-/circSfl-FLAG encoded a T7 promoter followed by a 5' external guide sequence (EGS), Anabaena group I intron, E30, circRNA UTR, circRNA CDS, link to a 3X FLAG sequence via Ser-Ser linker, E15, 3' EGS, EcoRV cleavage site. For generation of circMbl-/circSfl-Nat the CJ was placed in the CDS and synonymous mutations were introduced to generate the required eACA structure [2]. The IGS was adapted accordingly; other than that, the Nat and NLuc constructs shared the same architecture as the FLAG constructs. Both, the circRNA FLAG and Nat circularization constructs were synthesized by IDT and provided pre-cloned into pUCIDT-Kan. The circRNA NLuc constructs were based on previous vectors [2]; here, the CJ was placed in the NLuc ORF and circMbl/Sfl sequences, CrPV or CSFV IRESes (synthesized by IDT or Genewiz, respectively), human β-globin 5' UTR, or no extra sequences (circNLuc) were then inserted between the C- and N-terminal fragments of the NLuc ORF (i.e. between the stop and start codon, respectively) via PCR and Gibson assembly. Gly-Ser-Gly linkers were added to connect the circRNA CDS to the NLuc ORF.

For recombinant production of human eIF4B and eIF4G (residues 165-1599), cDNA sequences were amplified by PCR from a cDNA library and then cloned in pFastBac1 containing an N-terminal Twin Strep tag followed by a PreScission protease cleavage site.

Gene blocks containing the coding sequences for SARS-CoV2 Nsp1 and human YBOX1, PA2G4 and DHX29, codon-optimized for expression in *E. coli*, were synthesized by IDT, then cloned into corresponding expression vectors: Nsp1: pET28a (N-terminal tags: MBP - TEV protease cleavage site - $His_6$ - Twin Strep tag - PreScission protease cleavage site); YBOX1, PA2G4: pCA528-SUMO (N-terminal $His_6$ - SUMO tag); DHX29: pFastBac1 (N-terminal tags: Twin Strep tag – FLAG tag – PreScission protease cleavage site).

All cloning was done using standard methods, including Q5 (NEB, cat. #E0555) driven PCR, Gibson assembly using HiFi DNA assembly mix (NEB, cat. #E2621), QuickChange mutagenesis, and DNA ligation employing 5' phosphorylated primers (IDT) and T4 DNA Ligase (NEB, M0202) for creation of deletion constructs. *E. coli* TOP10 was used as cloning strain.

#### circRNA preparation

Production of circRNAs is based on Du et al.[2] For template preparation of circMbl/Sfl Nat and FLAG constructs, *E. coli* TOP10 cells transformed with plasmids encoding circRNA precursors under the control of a T7 promoter were grown in 120 ml LB medium o.n. cultures. Plasmids were purified using a Maxiprep kit (Qiagen, cat. #12963) and subsequently linearized by EcoRV-HF (NEB, cat. #R3195) digestion. The templates were then extracted using a mix of phenol:chlorophorm:isoamyl alcohol (P:C:I, Sigma, cat. #77617), followed by EtOH precipitation and subsequently re-dissolved in diethyl pyrocarbonate (DEPC) treated $H_2O$. Templates for all circRNA NLuc constructs were generated via PCR using Q5 DNA Polymerase (NEB, cat. #E0555) from plasmids encoding the circularization constructs; templates were subsequently isolated using a PCR purification kit (Qiagen, cat. #28104). *In vitro* transcription was then performed in reactions containing 80 mM HEPES/KOH (pH 7.5), 2 mM spermidine, 40 mM DTT, 14 mM $MgCl_2$, each 6 mM GTP, ATP, CTP and UTP (all from Sigma), 0.2 U/µl RNasin (Promega; cat. #N2511), 0.014 µg/µl home-made T7 RNA polymerase, 1 µg/ml yeast pyrophosphatase (Roche, cat. #10108987001), and ~50 ng/µl linearized plasmid or 15-20 ng/µl PCR product as templates in DEPC-$H_2O$. Reactions were incubated for 6 h or o.n. at 37 °C, then templates were digested by adding 30 U RNAse-free DNAse (NEB, cat. #M0303) per 1 ml reaction and further incubation for 30 min at 37 °C. RNAs were then collected by LiCl precipitation and subsequently resuspended in DEPC-$H_2O$. Prior to circularization, the precursor RNAs were refolded (5 min at 98 °C, then 3 min on ice). Circularization reactions containing 0.5 µg/µl precursor RNA in 50 mM Tris/HCl (pH 7.4), 10 mM $MgCl_2$, 1 mM DTT, 2 mM GTP were then set up and incubated for 30 min at 55 °C. The circularized RNAs were collected by LiCl or EtOH precipitation and subsequently resuspended at approximately 2 µg/µl in DEPC-$H_2O$. CircRNAs were then purified by size exclusion chromatography (SEC) using a 4.6x30 mm SRT SEC-2000 column (Sepax, cat. #215980P-4630) run in 10 mM Tris/HCl, 1 mM EDTA (pH 6.5 at room temperature) in a cold room. In each run, 50 µl of RNA (up to ~100 µg) were injected and 70 µl fractions were collected, while keeping UV detection off to avoid potential nicking. Fractions were analyzed by native agarose gel electrophoresis, circRNA containing fractions combined and concentrated by EtOH or isopropanol precipitation. Typically, removal of most unreacted precursor RNA and intron was achieved in two – three consecutive runs. After the final SEC run, nicked circRNAs were removed by RNAse R digestion (Abcam, cat. #ab286929) in reactions containing ~0.5 µg/µl RNA, and ~3-5 U RNase R/10 µg RNA in RNase R buffer. After incubation for 25 – 30 min at 37 °C, the circRNAs were purified using Zymo IIICG spin columns (cat. #C1006), eluted in DEPC-$H_2O$ at >500 ng/µl, aliquoted and then shock-frozen and stored at -70 °C.

**Preparation of linear mRNAs**

For preparation of VHP-β mRNA, plasmid pSP64_3X-FLAG-VHP-b-ΔTM (encoding Sp6 promoter, 65 nt 5' UTR, 3X-FLAG, VHP-β ORF, 50 nt 3' UTR; [3]) was used as template for

PCR with Q5 DNA polymerase (NEB, cat. #E0555). The PCR product was purified (Qiagen, QIAquick PCR purification kit, cat. #28104), then used at 25 ng/µl in an *in vitro* transcription reaction containing 80 mM HEPES/KOH (pH 7.5), 32 mM $MgCl_2$, 2 mM spermidine, 40 mM DTT, 4 mM of each GTP, ATP, UTP and CTP (all Sigma), 0.2 U/µl RNasin (Promega; cat. #N2511), 0.2 U/µl SP6 RNA polymerase (NEB, cat. #M0207), and 2 µg/ml yeast pyrophosphatase (Roche, cat. #10108987001) in DEPC-$H_2O$. *In vitro* transcription was carried out o.n. at 37 °C, then RNAse-free DNAse I (20 U/ml reaction; NEB, cat. #M0303) was added, and reactions were further incubated at 37 °C for 45 min. The RNA was then precipitated using LiCl, resuspended in DEPC-$H_2O$ and subsequently capped using the Vaccinia capping system (NEB, cat. #M2080). The capped RNA was precipitated using LiCl and subsequently purified by preparative urea PAGE (7 M urea, 5 % polyacrylamide gel) and electroelution using an EluTrap device. The purified RNA was then polyadenylated using *E. coli* polyA polymerase (NEB, cat. #2076), subsequently purified using an RNA Clean & Concentrator 100 kit (Zymo, cat. #D4039) and finally aliquoted, flash frozen and stored at -70 °C.

NLuc mRNA containing a β-globin 5' UTR (β-gl NLuc) was transcribed from a T7 promotor using a linearized plasmid as the template. Template preparation and *in vitro* transcription were carried out as described for preparation of circRNA precursors. The raw RNA was then capped using the Vaccinia capping system (NEB, cat. #M2080) according to manufacturer's instructions, then purified using an RNA Clean & Concentrator 100 kit (Zymo, cat. #D4039), and subsequently polyadenylated using *E. coli* polyA polymerase (NEB, cat. #2076). The RNA was again purified by using Zymo IIICG spin column (cat. #C1006), finally aliquoted, shock frozen stored at -70 °C.

**Urea agarose gel electrophoresis**

Urea agarose solutions containing 6 M urea, 1.5 % agarose in TBE were kept at RT until use, solutions containing >1.5 % agarose were prepared fresh. Gels were casted using either empty 1.5 mm mini gel cassettes (Invitrogen, cat. #NC2015) or BioRad 1.5 mm glass plates that were modified with a horizontal spacer at the bottom to contain the gel between the gels during the run, and the agarose allowed to solidify o.n. at 4 °C. Gels were usually run for 20 – 25 min at 17 – 20 W in a cold room using TBE buffer, then chilled on ice for 15 min to re-solidify any molten agarose and finally stained using SYBR Green II (Invitrogen, cat. #S7564) or SYBR Gold (Invitrogen, cat. #S11494) stain in TBE.

**Production of proteins, ribosomal subunits and tRNA**

Human initiation factors eIF1, eIF1A, eIF5, eIF4E and eIF4A were expressed in *E. coli* BL21 (λDE3) as $His_6$-MBP-TEV fusions as described previously.[4] Proteins were purified according to ref.[4]. Briefly, cells were lysed using sonication, the raw extract applied to Ni affinity chromatography, followed by cleavage of the tag with TEV protease o.n., ion exchange (MonoS for eIF1, eIF1A and eIF4E, and MonoQ for eIF5 and eIF4A) and SEC using 20 mM HEPES (pH 7.5), 200 mM KCl, 10 % glycerol, 1 mM DTT as the final buffer. For preparation of human PABP, we used a $His_6$-TEV construct described in ref.[5] and purified the protein according to established purification methods described.[4]

Endogenous human eIF2 and eIF3 were purified from HeLa cells as described earlier.[4,6]

Human eIF4B and eIF4G (165-1599) contained a Twin Strep tag followed by a PreScission cleavage site at the N-terminus and were expressed in Sf9 insect cells following standard procedures. Cells containing the overexpressed factors were resuspended in lysis buffer (20 mM HEPES (pH 7.5), 300 mM KCl, 10 % glycerol, 1 mM TCEP, 0.1 % Triton X-100, supplemented with one cOmplete protease inhibitor tablet (EDTA-free; Roche, cat #11873580001) per 50 ml), then lysed by sonication. The lysate was cleared by centrifugation, then applied to a 5 ml StrepTrap HP column affinity column (Cytiva, cat. #28907547). The proteins were eluted with lysis buffer supplemented with 2.5 mM D-desthiobiotin and the tag was then cleaved off using PreScission protease overnight at 4 °C. The sample was then applied to a 5 ml HiTrap Heparin HP column (Cytiva, cat. #17040703) and the proteins eluted using a 100-500 mM KCl gradient in 50 mM HEPES (pH 7.5), 10 % glycerol, 1 mM DTT. The factors were then concentrated and subjected to a Superose6 10/300 GL SEC column equilibrated in SEC buffer (50 mM HEPES (pH 7.5), 150 mM KCl, 10 % glycerol, 1 mM DTT). Fractions containing the pure proteins were then concentrated, the samples aliquoted, flash-frozen and stored at -80 °C.

Human 40S and 60S ribosomal subunits were purified from HEK293F cells collected at exponential phase. Cells were resuspended in lysis buffer (50 mM HEPES (pH 7.5), 10 mM KCl, 1.5 mM $MgCl_2$, 5 mM DTT, 1 mM EDTA, 1 mM EGTA, supplemented with 1 tablet of cOmplete protease inhibitor (EDTA-free; Roche, cat #11873580001) per 50 ml), lysed by nitrogen cavitation, and cell debris and the mitochondrial fraction was subsequently removed by consecutive centrifugation at 800 x g, 1,000 x g and 10,000 x g for each 15 min at 4 °C, transferring the supernatant to a new tube between centrifugations. The cell lysate was further clarified by centrifugation at 20,000 x g for 30 min at 4 °C. Crude ribosomes were pelleted by applying 50 ml of lysate on a 10 ml sucrose cushion (50 mM HEPES (pH 7.5), 150 mM KCl, 10 mM $MgCl_2$, 5 mM DTT, 1 mM EDTA, 1 mM EGTA, 8.5 % D-mannitol, 1 M sucrose, supplemented with one tablet of cOmplete protease inhibitor (EDTA-free; Roche, cat

#11873580001) per 50 ml) and centrifugation at 230 x g for 16 h at 4 °C in a Ti45 rotor (Beckman Coulter). After centrifugation, pellet were resuspended in 3 ml of gradient buffer 50 mM HEPES (pH 7.5), 150 mM KCl, 10 mM $MgCl_2$, 1 mM DTT) and aliquots of 2 ml were flash frozen in liquid nitrogen and stored at -80 °C. Aliquots of crude ribosomes (~300 $A_{260}$ units) were separated on 40 ml 15-40% (w/v) sucrose gradients in gradient buffer at 80,000 x g for 16 h at 4 °C. Gradients were fractionated using a Piston Gradient Fractionator (BIOCOMP Instruments); 80S peak fractions were collected and ribosomes subsequently pelleted by centrifugation at 235,000 x g for 16 h at 4 C. After removing the supernatant, each pellet was resuspended in 400 μl of separation buffer (50 mM HEPES (pH 7.5), 500 mM KCl, 3 mM $MgCl_2$, 1 mM DTT) to split the ribosome in 40S/60S subunits. Samples from six gradients were combined and puromycin dihydrochloride was added to 2 mM and the mixture was incubated at 32 °C for 10 min to release nascent chains from 60S-bound aminoacyl tRNAs. Ribosome subunits were separated on six 40 ml 10-35% (w/v) sucrose gradients in separation buffer by centrifugation at 110,000 x g for 16 h at 4 °C using a SW-28 rotor (Beckman Coulter). Gradients were fractionated using a Piston Gradient Fractionator (BIOCOMP Instruments), fractions corresponding to 40S and 60 S subunits were combined separately, and ribosomes were pelleted by centrifugation at 235,000 x g for 16 h at 4 C in a Ti 45 rotor (Beckmann Coulter). Supernatants were removed and ribosomal pellets were resuspended in storage buffer (50 mM HEPES (pH 7.5), 50 mM KCl, 5 mM $MgCl_2$, 1 mM DTT) at concentrations >4 μM, aliquoted, flash frozen in liquid nitrogen and stored at -80 °C.

Native yeast $tRNA_i^{Met}$ was overexpressed in *S. cerevisiae* harbouring the high copy plasmid pKC35 [7]. 24 L of yeast culture was grown to OD600 ~4. Cells were collected by centrifugation and resuspended in lysis buffer (1 mM Tris-HCl (pH 7.5) 10 mM $Mg(OAc)_2$; ~10 ml per 1 L of cell culture). The cell suspension was passed twice through a cell disruptor at 35,000 psi. Debris was then removed by centrifugation at 20,000 rpm, 4 °C for 30 min (JA 25.5 rotor; Beckmann Coulter). Total RNA was extracted from the supernatant by addition of an equal volume of citric acid-equilibrated phenol (pH 4.3; Sigma, cat. #P4682) and stirring for 30 min. The aqueous phase was transferred to a clean tube and precipitated with 3 volumes of ethanol at -20 °C. After centrifugation at 20,000 rpm, 4 °C for 20 min, the supernatant was removed, and the pellet was resuspended in 200 ml of 1 M NaCl. The suspension was centrifuged, the supernatant then transferred to clean tubes and precipitated with 3 volumes of ethanol at -20 °C. The pellet was again collected by centrifugation and then resuspended in 100 ml 1.5 M Tris-HCl (pH 8.8) and incubated in a 37 °C water bath for up to 2 h to hydrolyze all tRNAs-amino acyl ester bonds. RNA was then precipitated with 3 volumes of ethanol at -20 °C. After centrifugation, the pellet was resuspended in 50 ml Q-Sepharose buffer (20 mM Tris-HCl (pH 7.5), 8 mM $MgCl_2$, 200 mM NaCl, 0.1 mM EDTA) and the solution filter through a 0.45 μm syringe filter. RNA was subsequently applied to an in-house packed 120 ml Q-sepharose column. The column was washed with buffer until a stable base line was established, and the RNA then eluted with a 0-1 M NaCl gradient in Q-sepharose buffer over 750 ml; 4 ml fractions were collected. Fractions from the left shoulder of the first elution peak were combined and RNA precipitated with 3 volumes of ethanol at -20 °C. After centrifugation the pellet was resuspended in ~120 ml of 5PW buffer A (10 mM $NH_4OAc$ (pH 6.3), 1.7 M $(NH_4)_2SO_4$). About ⅓ to ¼ of the sample was loaded at a time on a 60 ml TSKgel Phenyl 5PW column (TOSOH, cat. #07656) equilibrated in 5PW buffer A. The column was washed with of 5PW buffer A until a stable base line was established and RNA was then eluted with 0-50 % gradient of 5PW buffer A to 5PW buffer B (10 mM $NH_4OAc$ (pH 6.3)) over 120 ml; 2 ml fractions were collected. Fractions from the first large elution peak (conductivity ~187-184 mS/cm) of each run were combined and RNA precipitated with 3 volumes of ethanol at -20 °C. After centrifugation the pellet was resuspended in ~12 ml of C18 buffer A (20 mM Tris-acetate (pH 6.0), 20 mM $Mg(OAc)_2$, 400 mM NaCl). About ¼ of the sample was loaded at a time on a SUPELCOSIL C18 column (Supelco, cat. #58355-U) equilibrated in C18 buffer A. The column was washed with C18 buffer A and RNA was then eluted with a 0-50 % gradient of C18 buffer A to C18 buffer B (20 mM Tris-acetate (pH 6.0), 20 mM $Mg(OAc)_2$, 400 mM NaCl, 60% methanol) over 45 ml; 0.5 ml fractions were collected. Fractions of the largest peak from each run were combined and the RNA then concentrated and buffer exchanged into acylation buffer (40 mM Tris (pH 7.6), 10 mM $Mg(OAc)_2$, 150 mM KCl, 10 mM DTT, 5 % DMSO). The tRNA was subsequently amino acylated as described in [8]. Acylated Met-$tRNA_i^{Met}$ was isolated from non-acylated $tRNA_i^{Met}$ using a C18 column and running conditions as before. Fractions containing the charged tRNA were then combined, concentrated/buffer exchanged in storage buffer (10 mM $NH_4OAc$ (pH 5.0), 50 mM KCl), aliquoted and stored in liquid nitrogen.

MBP-TEV-His6-Twin-Strep-Nsp1 protein was expressed from pET28a in *E. coli* Rosetta (λDE3) cells grown in LB medium. Expression was induced with 1 mM IPTG at an OD600 of ~0.5, and carried out for 5 h at 37 °C. For protein purification, cells were suspended in lysis buffer (20 mM Tris/HCl (pH 7.4), 500 mM NaCl, 20 mM imidazole, 1 mM TCEP), lysed by sonication and the raw extract subsequently applied to a 10 ml Ni-NTA agarose gravity flow column (Qiagen, cat. #30210). The protein was eluted using lysis buffer containing 500 mM imidazole and 150 mM NaCl. The tag was then cleaved off over-night at 4 °C using PreScission protease and the sample then further purified via ion exchange chromatography using a 5 ml HiTrap Q HP column (Cytiva, cat. #17115401), followed by another round of Ni-NTA affinity chromatography. The flow-through of the Ni column was collected, the protein then buffer exchanged into storage buffer (20 mM HEPES (pH 7.4), 150 mM NaCl, 1 mM TCEP) using a spin concentrator, concentrated, aliquoted, flash-frozen and stored at -80 °C.

N-terminally $His_6$ tagged *E. coli* RelE protein was purified as described in [9].

$His_6$-SUMO-YBOX1 was expressed from pCA528 in *E. coli* Rosetta (λDE3) cells grown in LB medium. Expression was induced with 0.2 mM IPTG at an OD600 of ~0.6-0.8, and carried out o.n. at 16 °C. For protein purification, cells were suspended in lysis buffer (40 mM Tris (pH 7.4), 500 mM NaCl, 5 % glycerol), lysed using sonication, and the lysate cleared by centrifugation. Nucleic acids were subsequently precipitated by the addition of 1/10 volume of 10 % polyethyleneimine (PEI) and centrifugation. The supernatant was then supplemented with 1.8 M $(NH_4)_2SO_4$ to precipitate the proteins from the PEI containing solution. The pellet was again collected by centrifugation, then redissolved in lysis buffer and applied to a 10 ml Ni-NTA agarose gravity flow column (Qiagen, cat. #30210). The target protein was eluted using a step gradient of elution buffer (40 mM Tris (pH 7.4), 150 mM NaCl, 5 % glycerol) containing 50, 100 or 300 mM imidazole, respectively. The tag was then cleaved using *S. cerevisiae* SUMO protease (Ulp1) over-night at 4 °C during dialysis in dialysis buffer (40 mM Tris (pH 7.4), 150 mM KCl). The sample was then re-applied to the 10 ml Ni-NTA column to remove uncleaved $His_6$-SUMO-YBOX fusion protein and protease, and the FT collected. The sample was then concentrated and applied to a Superdex75 10/600 GL SEC column (Cytiva, cat. #17517401) equilibrated in SEC buffer (40 mM Tris (pH 7.4), 200 mM KCl, 5 % glycerol, 2 mM DTT). Fractions containing the target protein were combined, concentrated, aliquoted and flash-frozen, then stored at -80 °C.

$His_6$-SUMO-P2AG4 was expressed as described for $His_6$-SUMO-YBOX1. Cells were resuspended in lysis buffer (40 mM Tris (pH 7.4), 500 mM NaCl, 5 % glycerol), lysed via sonication, and the lysate cleared by centrifugation. The raw-extract was then applied to a 10 ml Ni-NTA agarose gravity flow column (Qiagen, cat. #30210), and the target protein was eluted using a step gradient of elution buffer (40 mM Tris (pH 7.4), 150 mM NaCl, 5 % glycerol) containing 40, 70, 150 or 300 mM imidazole, respectively. The tag was then cleaved using *S. cerevisiae* SUMO protease over-night at 4 °C during dialysis in dialysis buffer (40 mM Tris (pH 7.4), 50 mM KCl). The sample was then re-applied to the 10 ml Ni-NTA column, the FT containing the target protein without tag collected and then applied to a MonoQ 10/100 GL column (Cytiva, cat. #17516701) to remove nucleic acids; elution was performed using a 50-500 mM KCl gradient. Fractions containing P2AG4 were combined, concentrated and the applied to a Superose 6 10/300 GL SEC column (Cytiva, cat. #17517201) equilibrated in SEC buffer (40 mM Tris (pH 7.4), 200 mM KCl, 5 % glycerol, 2 mM DTT). Fractions containing pure target protein were combined, concentrated, aliquoted, flash-frozen and stored at -80 °C.

Twin Strep-FLAG-PreScission cleavage site-tagged human DHX29 was expressed in Sf9 insect cells according to standard procedures. Cells containing the over-expressed factor were resuspended in lysis buffer (40 mM Tris (pH 6.0), 500 mM NaCl, 5 % glycerol, 0.5 mM EDTA, 0.1 % Triton X-100, supplemented with one cOmplete protease inhibitor tablet (EDTA-free; Roche, cat #11873580001) per 50 ml), then lysed by sonication. The lysate was cleared by centrifugation and the supernatant then applied to a 5 ml StrepTrap HP column (Cytiva, cat. #28907547). The factor was eluted using lysis buffer supplemented with 2.5 M D-desthiobiotin and fractions containing the target protein were concentrated and applied to a Superose 6 10/300 GL SEC column (Cytiva, cat. #17517201) equilibrated in SEC buffer (40 mM Tris (pH 6.0), 200 mM KCl, 5 % glycerol, 1 mM DTT). Fractions containing the pure factor were then concentrated, aliquoted, flash-frozen and stored at -80 °C.

##### ***In vitro*** **translation**

RRL *in vitro* translation assays were based on [10]. Briefly, 10 µl reactions consisting of ½ volume of nuclease treated RRL with supplements [10] was mixed with amino acid mixture minus Methionine (10 µM final concentration; Promega, L9961), RNasin (1 U/µl; Promega, cat. #N2511) and EasyTag L-$^{35}$S-Methionine (0.5 µCi/µl final; revvity, cat. #NEG709A001MC). Positive control mRNA (capped and polyadenylated VHP-β) or circRNAs were added at 100 nM and translation was conducted for 30 min (positive control mRNA) or 45 min (circRNAs), respectively, at 32 °C using a water bath. Aliquots were then analyzed by SDS-PAGE, gels were stained, subsequently dried, and bands visualized by autoradiography using an Imaging plate (Amersham BAS-IP SR 2025, cat. #28956478) or X-Ray film (Photon Imaging Systems, cat. #P10M000266A). Autoradiographs were analyzed using ImageJ v. 1.53k.[11]

*In vitro* translation experiments involving HeLa lysate were conducted using the One-Step Human IVT kit (Thermo, cat. #88881) according to manufacturer's instructions. Briefly, 20 µl reactions were assembled using HeLa lysate, reaction mix and accessory proteins; circRNAs and VHP-β mRNA were added to 70 nM and translation was carried out for 1.5 h at 32 °C. Samples were then analyzed by anti-FLAG Western blotting (primary antibody: mouse anti-FLAG antibody (Sigma, cat. #F1804) at 1:1,000 dilution; secondary antibody: Alexa Fluor Plus 800-conjugated goat anti-mouse antibody (Thermo, cat. #A32730) at 1:20,000 dilution).

##### **RNA transfection and NLuc activity measurements**

RNA transfection was performed using a reverse transfection protocol. HEK293T and HeLa S3 cells were maintained in adherent culture at 37 °C, 5 % $CO_2$, using DMEM containing 10 % FBS. *Drosophila* S2 cells were grown as semi-adherent cells at 27 °C in Schneider's Drosophila medium (Gibco, cat. #21720-024) containing 10 % heat-inactivated FBS (Gibco, cat. #10270106).

Prior to transfection, RNAs were refolded in DEPC-$H_2O$ (3 min at 65 °C, then 5 min on ice) and then diluted with either serum-free DMEM or Schneider's Drosophila medium to 0.02 pmol/µl. Lipofectamine MessengerMAX Transfection reagent (Invitrogen, cat. #LMRNA003) was diluted in the corresponding medium (0.3 µl in 5 µl of medium), subsequently mixed with an equal volume of diluted RNA and incubated for 5 min at RT. The RNA:transfection reagent mix was then added to 96 well plates (10 µl per well, i.e. 0.1 pmol of RNA) followed by 100 µl of cells at a density of 100,000 cells/ml (i.e. 10,000 cells per well). The cultures were then incubated for 24 h. To quantify translation, cells were lysed using a buffer containing 25 mM Tris/HCl (pH 7.4), 150 mM NaCl, 10 mM $MgCl_2$, 1 mM EDTA, 2 % glycerol, 1 % Triton X-100 and NLuc activity was measured using the Nano-Glo Luciferase assay kit (Promega, cat. #N1110) according to the manufacturer's instructions and a Tecan Spark 10 M plate reader (integration time 1 s). Statistical analysis of NLuc activity measurements and preparation of bar graphs was done in GraphPad Prism v. 10.

**qRT-PCR**

RNA was reverse transfected (as described above) into 20,000 HEK293T cells per well in a 96-well plate and incubated for 24 h. Cells were gently washed in PBS before lysis in 100 µl TRIzol (Invitrogen, cat. #15596026) and total RNA isolated using TRIzol:Chloroform extraction followed by cleanup using the RNA Clean & Concentrator 5 Kit (Zymo Research, cat. # R1015). 100 ng RNA was reverse transcribed using SuperScript IV First-Strand Synthesis System with ezDNAse (Invitrogen, cat. # 18091150), and the resulting cDNA diluted with DEPC-$H_2O$ to 2 ng cDNA/µl (assuming 1:1 conversion of RNA to cDNA). 10 µl qRT-PCR reactions were assembled in (technical) triplicate for each gene in 384-well plates, composed of: 4 ng of diluted cDNA, 400 nM each of forward and reverse primers, and 1X iTaq Universal SYBR Green Supermix (Bio-Rad, cat. # 18091150). Reactions were run on the QuantStudio Pro 7 (Applied Biosystems) using the 'Standard' program setting, initial denaturation at 95°C for 30 s, and 40 cycles of: 95°C for 15 s, 60°C for 60 s. Cq data was exported and analysed in Microsoft Excel and GraphPad Prism, averaging technical triplicates and normalizing to GAPDH. The circNLuc primers were designed to flank the circularization junction (see Figure 1B), making amplification specific to cDNA derived from circRNAs vs. any traces of linear precursor.

#### Ribosomal toe-printing

Toe-printing assays were based on Pisarev et al.[12] Factors and RNAs were diluted in TP buffer (25 mM HEPES/KOH (pH 7.5), 100 mM KOAc, 2.5 mM $MgCl_2$, 0.25 mM spermidine, 1 mM DTT) and circRNAs were then refolded by heating for 1 min at 98 °C followed by immediate cooling for >3 min on ice. RNA activating factors, i.e. eIF4A (500 nM final concentration), eIF4G, eIF4B, eIF4E, DHX29, YBOX1, PA2G4, PABP (all at 250 nM final concentration) were added to the circRNAs (50 nM final concentration) along with 0.5 mM ATP (or ADP/AMP-PCP) and 0.5 U/µl RNasin (Promega; cat. #N2511). A 43S-PIC was assembled by mixing 40S subunits (100 nM), eIF1, eIF1A (both 500 nM), eIF2 (250 nM), yeast Met-$tRNA_i^{Met}$, eIF3 and eIF5 (all 200 nM) and incubating for 10 min at 30 °C. The circRNA:factor mix was then combined with 43S-PIC in a volume of 15 µl and complex formation was allowed for 15 min at 37 °C. After incubation, 5 µl of RT master mix (consisting of 1 µl 5' $^{32}$P labelled primer (~2 µM, labelled using T4 Polynucleotide kinase (NEB, cat. #M0201) and $\gamma$-$^{32}$P-ATP (Hartmann analytic, cat. #SRP-301)), 10 mM dNTPs mix, 0.875 µl 100 mM $MgCl_2$, 0.625 µl DEPC-treated-$H_2O$, and 1 µl 5X AMV RT buffer and 0.5 µl of 10 U/µl AMV RT (both Promega, cat. #M5108)) was added, and reverse transcription was conducted for 1 h at 37 °C in a water bath. Subsequently, the volume was adjusted to 100 µl with TES buffer (10 mM Tris/HCl (pH 8.0), 10 mM EDTA, 0.5 % (w/v) SDS) and cDNAs were extracted by P:C:I extraction, precipitated with EtOH and finally taken up in 5 µl of formamide/EDTA loading buffer. Aliquots of 2 µl were then run on 0.4 mm, 7 M urea, 6 % polyacrylamide sequencing gels next to sequencing ladders generated from the undigested plasmid used to prepare circRNA *in vitro* transcription templates, the corresponding toe-printing primer, $\alpha$-$^{32}$P-dATP (Hartmann analytic, cat. #SRP-203) and materials and instructions from the Sequenase Version 2.0 DNA Sequencing Kit (Affymetrix, cat. #70754). Gels were then dried and exposed to an Imaging Plate (FujiFilm BAS-IP MS 2040, cat. #28956474), that was imaged using a Typhoon Imager. The autoradiographs were then analyzed in ImageJ (v. 1.53k; [11]), where a background correction was applied.

#### Co-sedimentation assays

For interaction studies of RNAs with isolated ribosome subunits and 80S, components were mixed in 25 µl reactions buffered by 25 mM HEPES/KOH (pH 7.5), 100 mM KOAc, 2.5 mM Mg(OAc)2, 0.25 mM spermidine, 2 mM DTT. RNAs (circMbl-FLAG, circSfl-FLAG, precursor of circCrPV-NLuc and VHP-$\beta$ all at 200 nM) were incubated with or without 40S (200 nM) or 40S + 60S (200 nM and 260 nM), in the presence of 1 mM GTP and 0.5 U/µl RNasin (Promega; cat. #N2511) for 15 min at 30 °C. Then, 1.5 µl of 100 mM $Mg(OAc)_2$ was added (~8 mM final concentration) and the samples were incubated further for 5 min at 30 °C to allow formation of 80S ribosomes and stabilize potential complexes. The samples were then loaded to 1 ml of sucrose cushion (20 mM HEPES/KOH, 70 mM KOAc, 4 mM $Mg(OAc)_2$, 10 μM $Zn(OAc)_2$, 0.1 mM spermidine, 1 mM DTT, 3 % glycerol, 1 M sucrose) and spun for 1 h at 120,000 rpm, 4 °C in a TLA120.2 rotor using 11x34 mm polycarbonate tubes (Beckmann Coulter, cat. #343778). The supernatant was carefully recovered, and the pellets were resuspended in 100 µl of TE buffer. RNAs were then isolated from supernatant and pellets by P:C:I extraction followed by isopropanol precipitation. The RNA pellets were resuspended in each 10 µl of 1X formamide/EDTA RNA loading buffer, and aliquots were analyzed by urea agarose gel electrophoresis using 2.5 % agarose 6 M urea gel.

**RNA stability and nicking assays**

RRL *in vitro* transcription reaction containing circMbl- or circSfl-FLAG were assembled as described (see *In vitro* translation), except that $^{35}$S-Met was replaced with non-labeled amino acid. The reactions were incubated at 32 °C, for 0, 10, 20, 30 or 45 min, respectively. RNAs were then extracted using Trizol (Thermo, cat. #15596026) followed by isopropanol precipitation. Negative controls consisting of only RRL or pure circRNA were treated identically. Samples were then analyzed using 6 M urea, 3.5 % agarose (circMbl-FLAG) or 6 M urea, 2.5 % agarose gels (circSfl-FLAG).

Stability under toe-printing conditions was assessed by preparing samples containing either only circRNAs, or circRNAs together with eIF4G/4A/4B in presence or absence of the 43S-PIC as described above (see Ribosomal toe-printing for buffer, concentration of factors and 43S assembly) and incubating them for 15 min at 37 °C. The RNAs were then isolated using P:C:I extraction followed by cleanup using the RNA Clean & Concentrator 5 kit (Zymo, cat. #R1013), mixed with Formamide/EDTA RNA loading dye and analyzed on 6 M urea, 2 % agarose gels.

For nicking experiments, circRNAs (in DEPC-$H_2O$) were first refolded by incubation for 1 min at 98 °C followed by cooling for 1 min on ice, then mixed with nicking buffer (final concentration of components: 25 mM $NaHCO_3$, 1 mM EDTA). The samples were then boiled at 90 °C; at the indicated time points (see Figure S1), aliquots were removed from the heat immediately placed on ice to cool, and quenching buffer (final concentration of components: 50 mM Tris/acetate (pH 7.0), 1 mM $Mg(OAc)_2$) was added. Samples were then run on urea agarose gels and the ratios of circular vs nicked RNA was analyzed using ImageJ (v. 1.53k; [11]). The remaining samples were then immediately used for *in vitro* translation assays in RRL or toe-printing as described in the corresponding sections.

**Isolation of circRNA initiation complexes by sucrose gradient centrifugation**

For isolation of initiation complexes from RRL, 1 ml reactions were assembled that contained 500 µl of RRL mix [10], 20 µM L-Methionine, 0.2 U/µl RNAsin (Promega; cat. #N2511), 2 mM GMP-PCP (Sigma, cat. #M3509) and 2 mM $Mg(OAc)_2$, excluding the volume for RNA to be added. This mix was then incubated for 5 min at 32 °C. RNAs (circMbl-FLAG, circSfl-FLAG, linear VHP-β mRNA (capped and polyadenylated)) or $H_2O$ (negative control) were then added to a final concentration of 20 ng/µl and the reactions were further incubated for 30 min at 32 °C. Then 10 µl of 1 M $Mg(OAc)_2$ (10 mM final) was added to stabilize initiation complexes and the samples were layered on top of 10 – 30 % (w/v) sucrose gradients in 48S buffer (20 mM HEPES/KOH (pH 7.5), 70 mM KOAc, 4 mM $Mg(OAc)_2$, 10 µM $Zn(OAc)_2$, 0.1 mM spermidine, 1 mM DTT, 3 % glycerol; 500 µl per gradient). The samples were then centrifuged for 5.5 h at 40,000 rpm, 4 °C in a SW40 rotor and subsequently fractionated using a piston gradient fractionator device (BIOCOMP Instruments). The 40S/43S/48S containing fractions samples were combined, the ribosomal complexes subsequently pelleted by ultracentrifugation (3 h at 100,000 rpm, 4 °C in TLA100.3 rotor) and finally resuspended in 48S buffer. The samples were then used for protein identification by mass spectrometry.

**Mass spectrometry**

For protein identification in initiation complexes, samples obtained from sucrose gradients were digested with in solution with Trypsin and the resulting peptides analyzed by LC-MS/MS. Briefly, 25-30 µl samples were first treated with benzonase, then 4 mM DTT and 14 mM iodoacetamide were added, along with 5 µl of 0.2 µg/µl trypsin. Tryptic digestion was carried out at 37 °C overnight, then trifluoroacetic acid (TFA) was added to 0.5 % to acidify the samples. Peptides were subsequently cleaned up using an SCX disk, dried and finally resuspended in 3 % acetonitrile, 97 % HPLC water, 0.1 % TFA. The resulting samples were then analyzed by nano-scale capillary LC-MS/MS using an Ultimate U3000 HPLC (Thermo Fisher Scientific). A µ-precolumn cartridge (C18 Acclaim PepMap 100 (ThermoScientific Dionex)) was used to trap peptides prior to separation on a C18 Easy Spray column (Thermo Fisher Scientific). Peptides were eluted using a 8-40 % acetonitrile double slope gradient and then injected into in a QExactive HFX mass spectrometer (Thermo Scientific). Raw data were processed using Proteome Discoverer v3.1 (Thermo Fisher Scientific). A database search against the UniProt reference proteome for rabbit (*O. cuniculus*) was carried out for protein identification.

**CryoEM sample preparation and data collection**

Initiation complexes of circMbl were obtained by *in vitro* reconstitution. First, circMbl-FLAG was diluted to 1.5 µM using cryoEM buffer (40 mM HEPES/KOH (pH 7.5), 80 mM KOAc, 2.5

mM $Mg(OAc)_2$, 0.25 mM spermidine, 1 mM DTT), and then refolded by heating for 1 min at 98 °C, followed by cooling on ice for 3 min. The circRNA was subsequently mixed with eIF4G (300 nM), eIF4A (600 nM) and eIF4B (300 nM), along with 0.5 U/µl RNasin (Promega; cat. #N2511) and 0.5 mM AMP-PNP (Sigma, cat. #A2647). The circRNA:factor mix was prepared at 1.66x the final concentration and incubated for ~10 min on ice. A 43S-PIC was prepared by mixing 40S subunits (150 nM), eIF1 (450 nM), eIF1A (450 nM), eIF2 (300 nM), eIF3 (250 nM), Met-$tRNA_i^{Met}$ (250 nM), eIF5 (300 nM) and GMP-PCP (0.5 mM; Sigma, cat. #M3509) at 2.5x the final concentration of the components. The 43S sample was then incubated for 10 min at 30 °C to allow complex assembly. Then 9 µl of activated circRNA and 6 µl of 43S-PIC were then mixed and further incubated for 10 min at 37 °C.

The initiation complexes were stabilized by addition of 0.375 µl 100 mM $Mg(OAc)_2$ (5 mM $Mg^{2+}$ in final sample) and subsequently crosslinked by adding 1.67 µl of 25 mM BS3 (2.5 mM final concentration; Thermo, cat. #A39266) and incubation for 1 h on ice. The crosslinking reaction was then quenched by the addition of 1 µl 1 M Tris/HCl (pH 7.4) and further incubation for 15 min on ice.

For grid preparation, UltrAUfoil, R1.2/1.3 300 mesh grids (Quantifoil) were first washed with $H_2O$, then glow discharged for 5 min using an Edwards glow discharger at 0.1 torr and 30 mA. Subsequently, 3.5 µl of a 0.2 mg/ml graphene oxide suspension in $H_2O$ (Sigma, cat. #763705) was then applied to the top of the grids, blotted off after 1 min incubation at RT, and the grids were subsequently washed three times with 10 µl $H_2O$ on both sides. After drying (> 1 h) the grids were used for sample preparation using an Vitrobot Mark II (Thermo Fisher Scientific) equilibrated at 4 °C and 100 % relative humidity. 3.5 µl of sample was applied to the graphene oxide coated side and incubated for 30 s. Blotting was performed for 4 – 5 s at a blot force of -3 using a double layered Whatman 595 blotting paper before plunging into liquid ethane kept at -170 °C by a temperature-controlled cryostat.

Data were collected on a Titan Krios Mark III microscope (Thermo Fisher Scientific) at the LMB EM Facility, equipped with an energy filter and a Gatan K3 camera operated in counting mode, using EPU software in faster acquisition (AFIS) setting for acquisition. Images were collect using a defocus range of -1.2 to -3.0 µm, at a magnification of 105,000x, corresponding to a nominal pixel size of 0.826 Å/pix. The total dose was 40 e-/$Å^2$, distributed over 40 frames (i.e. 1 e-/$Å^2$ per frame).

**CryoEM data processing**

All datasets were processed with RELION v. 4.0 and 5.0 [13]. The raw micrographs were first motion-corrected using MotionCor2 [14], then the contrast transfer function (CTF) was estimated using CTFFIND v. 4.1 [15]. Micrographs showing a resolution worse than 5 Å were removed. Particles were picked using template matching of two classes of manually picked 48S complexes, then extracted using a 600 pixel box (495.6 Å) box, 4x down-scaled to 150 pixels (123.9 Å, 3.304 Å/pixel). An initial 3D classification was conducted to isolate ribosomal particles that contained both eIF3 and the TC, using a 20 Å low pass filtered version of a previously determined 48S scanning complex (EMD-11302, [16]) as a reference. The initial set of ribosomal particles was then re-extracted at the full pixel size and CTF refinement and polishing was performed. The AUG recognition and scanning complexes were separated by 3D classification without alignment, then a focused classification with signal subtraction on the TC of the scanning complex was performed to separate out particles lacking the TC. The full processing pipeline is depicted in Figure S4B. The final maps were then post-processed for model building and used for calculation of the local resolution using RELION's implementation. A 2D plot depicting the angular distributions of each final map was generated using the AngDist script (https://github.com/Guillawme/angdist).

**Model building, refinement and validation**

The model of a human 48S complex (PDB ID 8OZ0), containing 40S, eIF1A, eIF2$\alpha$ and $\gamma$, eIF3 (all subunits except eIF3j), eIF5, eIF4G, eIF4A, eIF4B and human Met-tRNA$_i^{Met}$ (without RNA modifications) was fitted into the postprocessed map of the circMbl-FLAG AUG recognition complex in UCSF Chimera v. 1.16 [17]. Further model building was then carried out using Coot v. 0.9.8.94 [18]. Previously identified modifications to the 18S rRNA as listed in [19] were added where missing. The map was blurred using B factor of 50 to enable fitting of regions with low local resolution, specifically eIF3 (except for eIFa and eIF3c), and eIF2$\alpha$ and $\gamma$. The initiator tRNA was replaced by a model obtained from a X-Ray crystal structure of native yeast initiator tRNA (PDB ID 1YFG), that contained all modifications. Since the density underneath the tRNA better matched eIF1, we replaced eIF5 with the eIF1 model from PDB ID 6ZMW. The RNA density in the channel was consistent with the sequence around the TIS (AAU ACG <u>AUG</u> GCC) and thus adapted correspondingly. The model was then inspected to correct any poorly fitted sites and minor mistakes of the previous models, and to add potential missing residues and ions.

The resulting coordinates were then used as a basis for modeling the circMbl-FLAG scanning complex. For this, the model was docked into the postprocessed map in Chimera, using the 40S body as the region for alignment. The 40S head is in a swiveled conformation compared to the AUG recognition complex, thus it was then docked separately into the correct conformation (this included 18S rRNA from G1211 – C1689 as well as the ribosomal proteins constituting the 40S head). The eIF2 chains were then replaced with those of PDB ID 6ZMW to reflect the presence of eIF2$\beta$. Any other changes were made in Coot, using a blurred map to fit any regions with low local resolution with locally restrained models. The RNA density in the 40S channel is much less defined than in the AUG recognition complex due to the lack of stabilizing interaction with the tRNA and eIF2$\alpha$. We therefore only model three nucleotides, based around the base making contacts with the C34 of $tRNA_i^{Met}$.

Refinement of all models was carried out using Phenix v. 1.21 [20]. A list of CIF files for modified RNA nucleotides was generated using phenix.elbow. The models were then refined using phenix.real_space_refine using the postprocessed maps and the corresponding models generated as described above. The non-bonded weight was set to 1000 and any rotamer outliers were fixed using option "outliers_or_poormap".

The final models were validated using phenix.validation_cryoem [20]. This program was also used to calculate the half-map and model-map FSC curves.

Models and maps for figures were depicted using UCSF ChimeraX v. 1.8 [21].

**Sequence analysis of circRNAs**

The GC content and sequence complexity of circMbl-FLAG and circSfl-FLAG were calculated in R, using the seqinr package and the sequence_complexity function (DUST algorithm) of the universalmotif package, respectively. For both calculations, a 25 nt sliding window was applied, circularity of the molecules was taken into account.

A list of translatable circRNAs (ribo-circRNAs) from *Drosophila* heads was obtained from table S2 in [22]. The circRNA sequences retrieved from the Genome browser using the corresponding genomic coordinates. Based on the annotation of the corresponding exons, circRNAs were classified as CDS-only, 5' UTR+CDS or others. CDS and UTR regions of the corresponding host mRNAs were then assigned and their lengths determined.

### Supplemental Figures and Tables

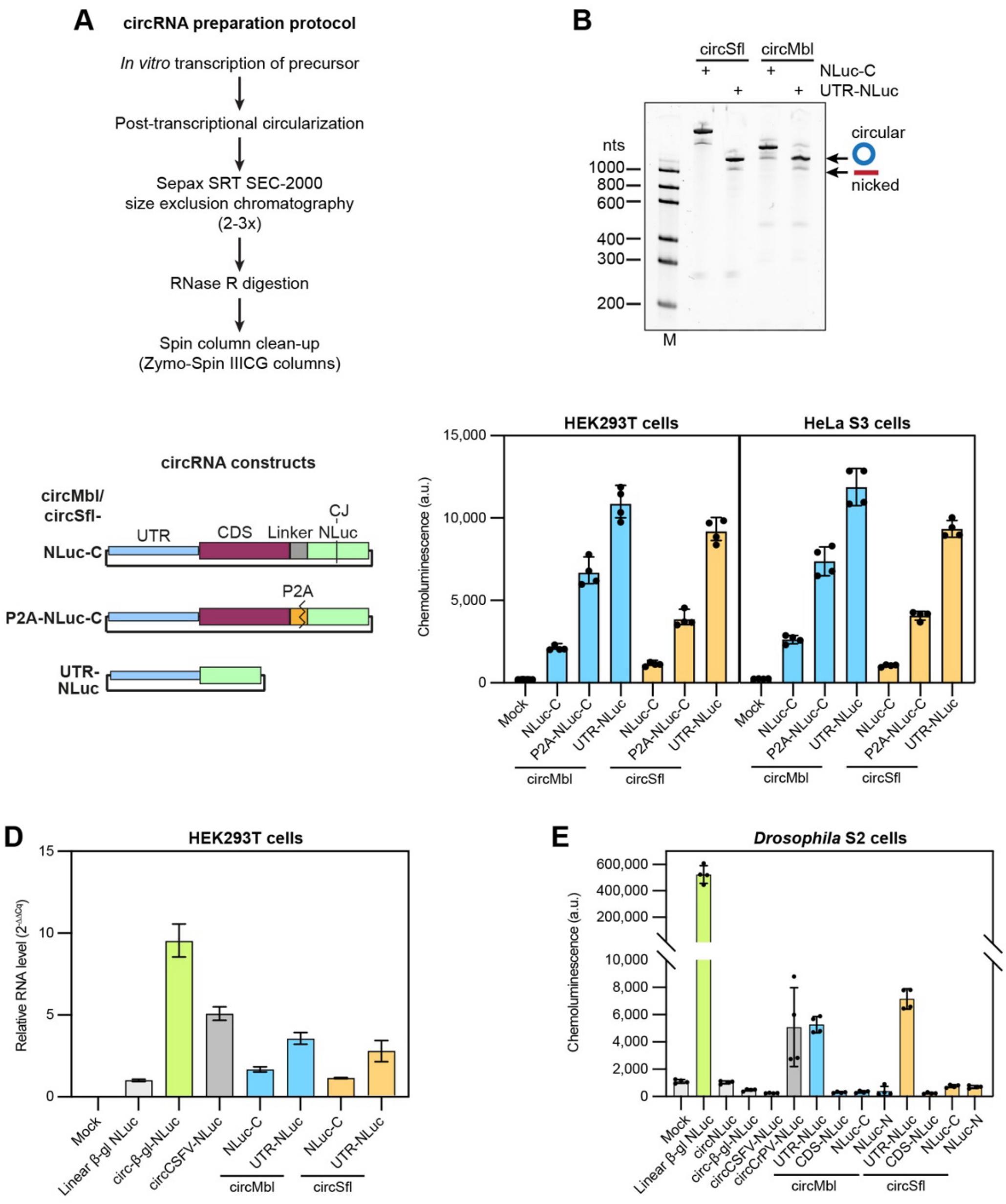


**Figure S1. Preparation and translation of naturally occurring circRNAs *in vitro* and in cells.**

(**A**) Outline of the circRNA preparation protocol.

(**B**) Urea agarose gel of selected circMbl/circSfl NLuc constructs. Bands representing the circular and linearized forms of the circRNAs are indicated.

(**C**) NLuc activity assays of circMbl/Sfl NLuc constructs in which the circRNA CDS-NLuc fusion protein has been severed co-translationally using a P2A sequence. Left: scheme of constructs. Right: NLuc activity measurements upon transfection of the constructs in HEK293T or HeLa S3 cells ($n$ = 4).

(**D**) Intracellular levels of mRNA/circRNA transfected into HEK293T assessed by qRT-PCR. The $2^{(-\Delta\Delta Cq)}$ value relative to linear β-globin mRNA (normalized to GAPDH) is given (*n* = 4 replicate wells on the same culture plate). Error bars indicate range derived from standard deviation of ΔΔCq.

(**E**) NLuc activity assays of *Drosophila* S2 cells transfected with circRNA NLuc constructs or a linear mRNA positive control (*n* = 4).

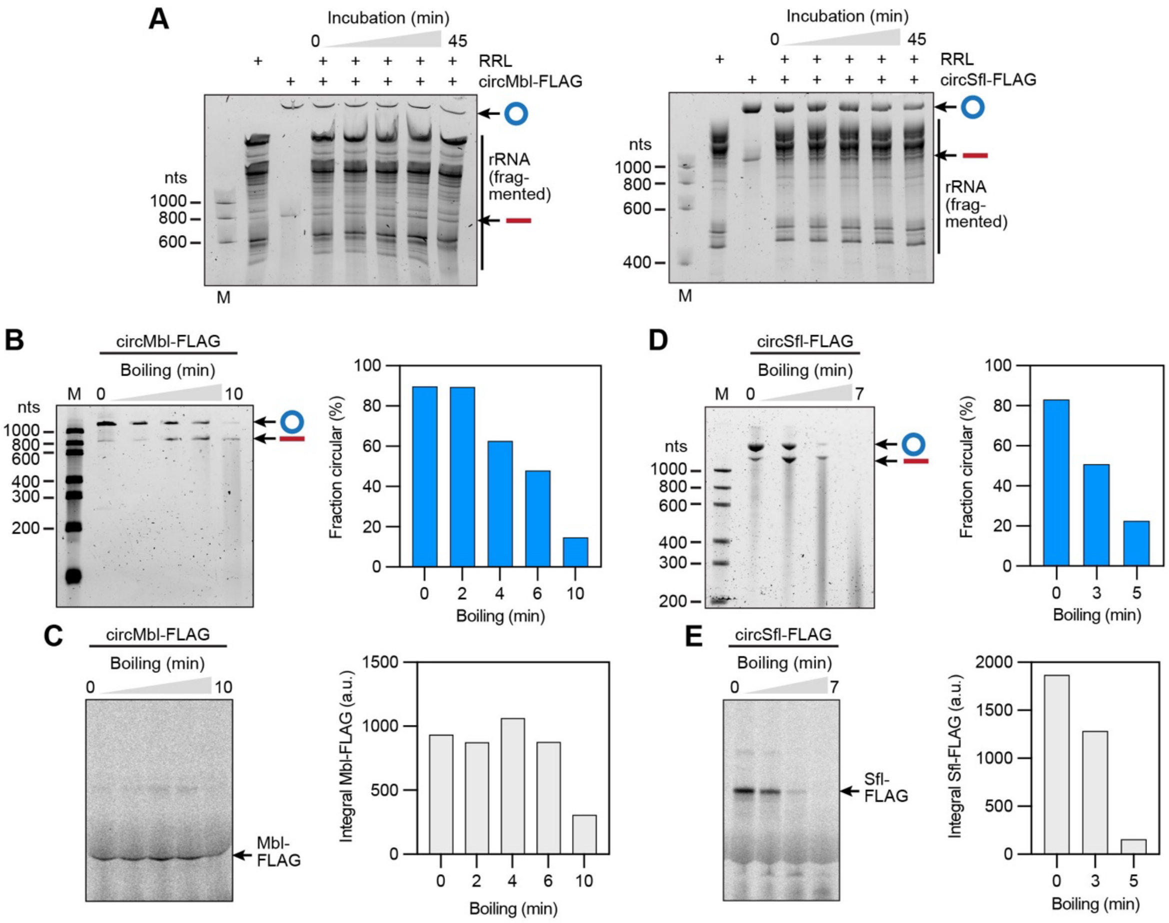


**Figure S2. CircRNAs are stable in cell lysates and nicking products do not substantially contribute to translational output under native conditions.**

(**A**) Urea agarose gels of circMbl-FLAG (left) or circSfl-FLAG (right) samples re-extracted after incubation in RRL over 45 min. Bands representing the circular and linearized forms of the circRNAs are indicated. Note that a nuclease-treated RRL was used for the experiment; therefore, rRNAs are partially fragmented.

(**B, C**) Translation of circMbl-FLAG nicking products in RRL. CircRNA was intentionally nicked by heating under alkaline conditions, nicking quenched after various incubation times and samples analyzed on a urea agarose gel (B, left), and used for *in vitro* translation assays in RRL followed by SDS-PAGE and autoradiography (C, left). Bands representing the circular and linearized forms of the circRNAs in (B) are indicated. The ratio of circular and nicked RNAs, or the amount of translated Mbl-FLAG quantified from the gels is depicted in the right panels in (B) or (C), respectively.

(**D, E**) Nicking experiment and RRL *in vitro* translation as in (B, C), but for circSfl-FLAG.

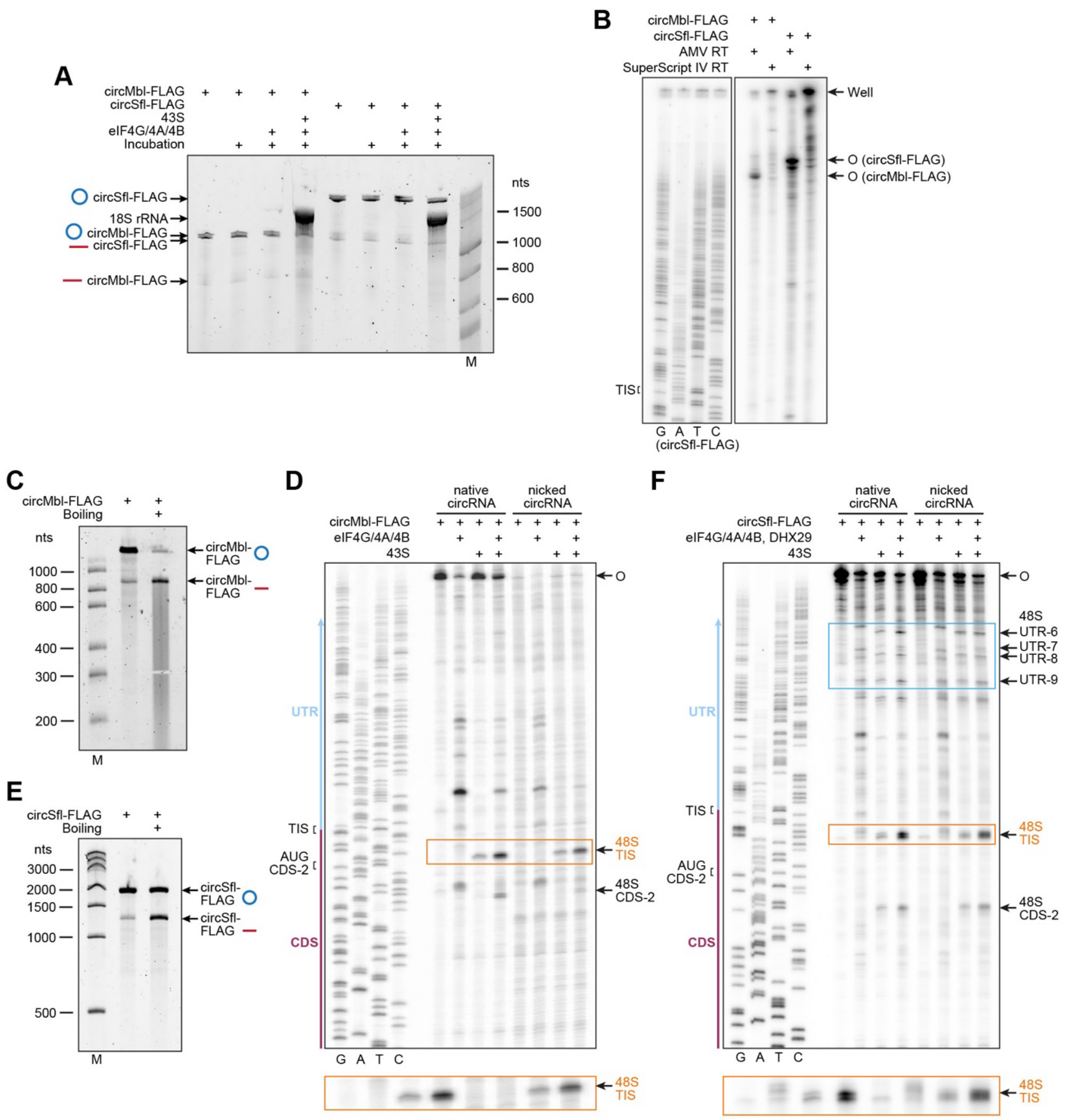


**Figure S3. Stability of circRNAs and contribution of nicking products to initiation complex formation under toe-printing conditions.**

(**A**) Urea agarose gels of circMbl-FLAG/ circSfl-FLAG samples re-extracted after incubation with the 43S-PIC and/or eIF4G/4A/4B under toe-printing conditions for 15 min. Bands representing the circular and linearized forms of the circRNAs are indicated.

(**B**) Autoradiograph of a sequencing gel showing a primer extension assay of circMbl-FLAG/circSfl-FLAG carried out using either AMV RT or SuperScript IV RT. AMV RT possesses an intrinsic RNase H activity that degrades circRNA in the hybrid with the cDNA, allowing synthesis of only one circle (indicated by O). SuperScript IV has little to no RNAse H activity allowing multiple rounds of cDNA synthesis to occur, resulting in a large product stuck in the wells.

(**C**) CircMbl-FLAG was intentionally nicked by heating under alkaline conditions and samples before and after heating analyzed on a urea agarose gel. Bands representing the circular and linearized forms of the circRNAs are indicated.

(**D**) Initiation complex formation on native and nicked circMbl-FLAG (see (C)) using a reconstituted system, as assayed by toe-printing. Note the absence of defined nicking products in the nicked sample, as well as the disappearance of full-length RT products (O). Furthermore, the amount of initiation complexes formed on the nicked sample is decreased compared to the intact sample, indicating that initiation complexes predominantly form on circular RNAs, but that some nicking products may also allow 48S complex formation.

(**E**) Nicking experiment done as in (C) but for circSfl-FLAG.

(**F**) Toe-printing assay done as in (D) but for circSfl-FLAG.

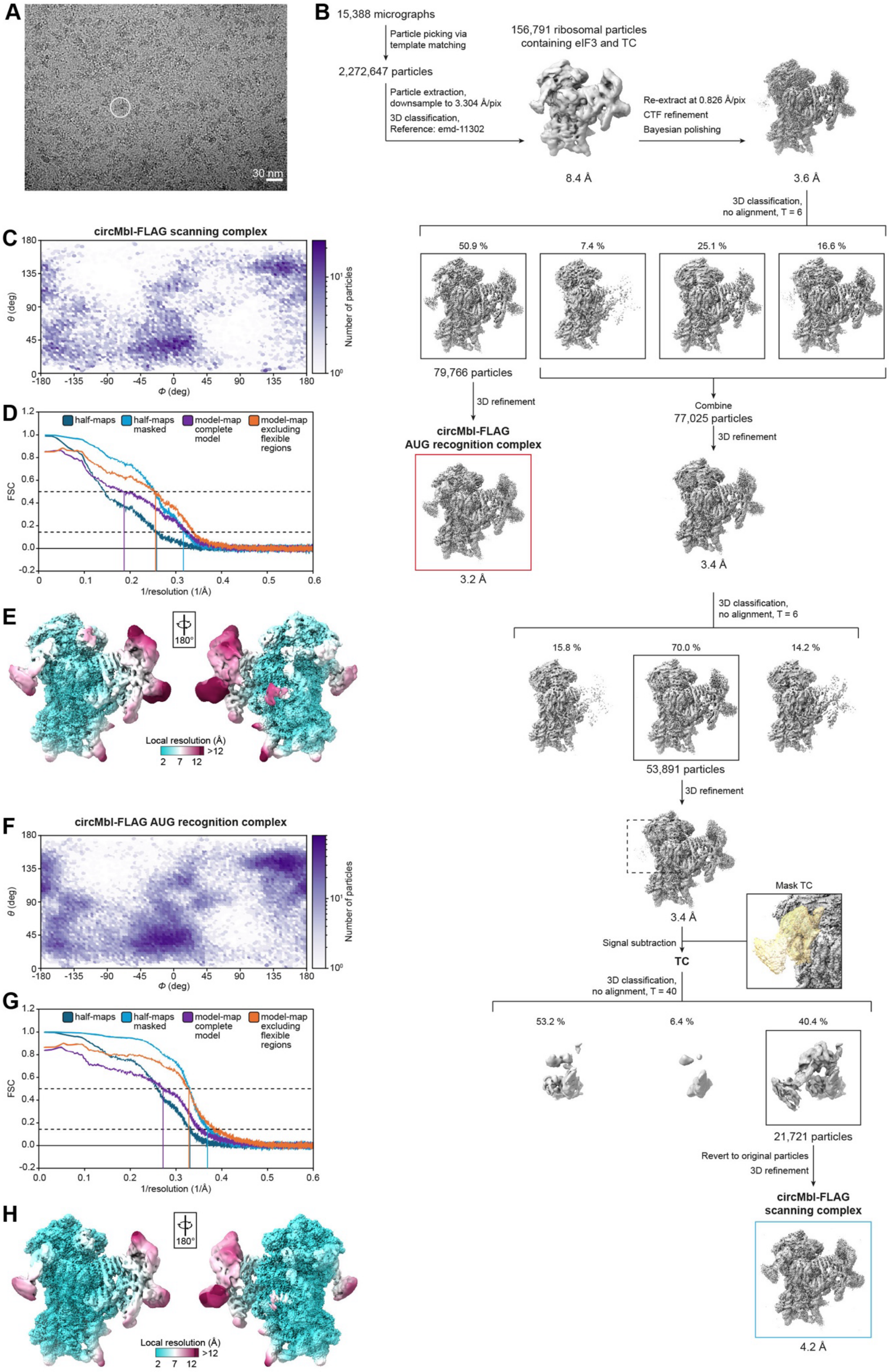
A
30 nm
B
15,388 micrographs
Particle picking via template matching
2,272,647 particles
Particle extraction, downsample to 3.304 Å/pix
3D classification, Reference: emd-11302
156,791 ribosomal particles containing eIF3 and TC
8.4 Å
Re-extract at 0.826 Å/pix
CTF refinement
Bayesian polishing
3.6 Å
3D classification, no alignment, T = 6
50.9 %
7.4 %
25.1 %
16.6 %
79,766 particles
3D refinement
circMbl-FLAG AUG recognition complex
3.2 Å
Combine
77,025 particles
3D refinement
3.4 Å
3D classification, no alignment, T = 6
15.8 %
70.0 %
14.2 %
53,891 particles
3D refinement
3.4 Å
Mask TC
Signal subtraction
TC
3D classification, no alignment, T = 40
53.2 %
6.4 %
40.4 %
21,721 particles
Revert to original particles
3D refinement
circMbl-FLAG scanning complex
4.2 Å
C
circMbl-FLAG scanning complex
θ (deg)
Φ (deg)
Number of particles
D
half-maps
half-maps masked
model-map complete model
model-map excluding flexible regions
FSC
1/resolution (1/Å)
E
180°
Local resolution (Å)
2 7 12 >12
F
circMbl-FLAG AUG recognition complex
θ (deg)
Φ (deg)
Number of particles
G
half-maps
half-maps masked
model-map complete model
model-map excluding flexible regions
FSC
1/resolution (1/Å)
H
180°
Local resolution (Å)
2 7 12 >12

607

**Figure S4. CryoEM data processing of the circMbl-FLAG initiation complex.**

(**A**) Representative micrograph of the circMbl-FLAG 48S sample. An exemplary 48S complex is highlighted.

(**B**) CryoEM data processing pipeline for the circMbl-FLAG initiation complex. The initial set of movies was processed using motion correction and CTF estimation, micrographs exhibiting a resolution worse than 5 Å were removed, resulting in 15,388 micrographs used for particle picking. RELION reported resolutions of the intermediate and final maps (after sharpening) are given, along with the number of underlying particles.

(**C**) Distribution of Euler angles of the final set of particles of the circMbl-FLAG scanning complex.

(**D**) Fourier Shell Correlation (FSC) curves of the circMbl-FLAG scanning complex. The half map FSCs (unmasked and using a solvent mask), and model-map FSC (unmasked, using either the complete model, or after exclusion of flexible regions [eIF2/eIF3], see Methods) are depicted, along with their intersections at FSC = 0.134 or 0.5.

(**E**) CryoEM map of the circMbl-FLAG scanning complex sharpened and colored according to local resolution.

(**F, G, H**) As in (C, D, E) but for the circMbl-FLAG AUG recognition complex.

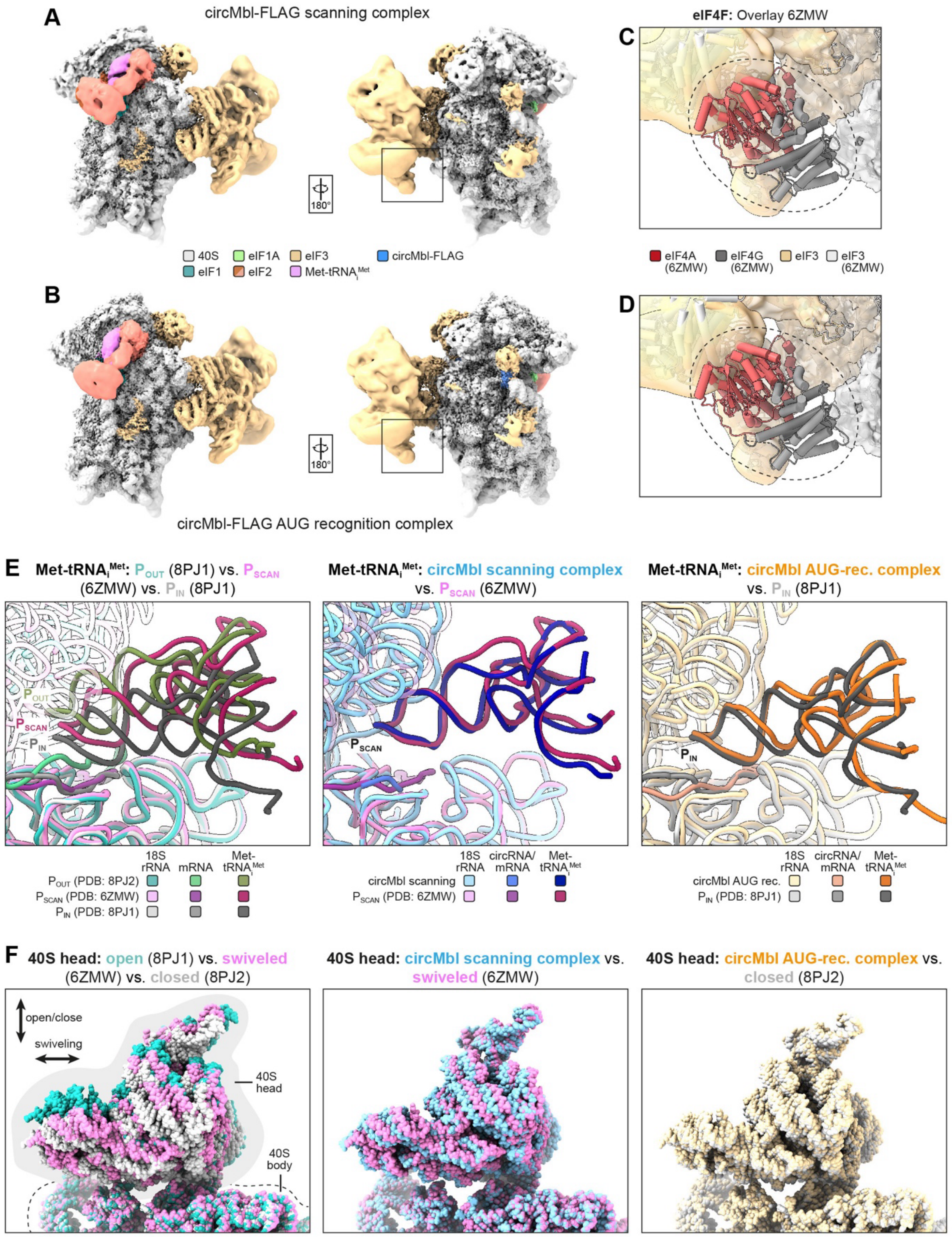


**Figure S5. Additional details of the structures of the circMbl-FLAG scanning and AUG recognition complexes.**

(**A, B**) Local resolution sharpened maps of the circMbl-FLAG scanning (A) and AUG recognition complexes (B), colours and relative orientations as indicated.

(**C, D**) Close-up view of the eIF4F interacting region on eIF3 boxed in (B, C, right). The model of the circMbl-FLAG initiation complexes was overlayed with the model of a previously determined scanning complex (PDB-ID 6ZMW). Maps are colored as in (A, B), models are colored as indicated. Note the absence of eIF4G/4A density in the circMbl-FLAG complexes. (**E, F**) Comparison of the position of the $tRNA_i^{Met}$ (E) or 40S head (F) between the circMbl-FLAG initiation complexes and previously determined structures of a scanning complex with $tRNA_i^{Met}$ in the $P_{OUT}$ position/40S head open (PDB ID 8PJ2), $tRNA_i^{Met}$ in $P_{SCAN}$ position/40S head swiveled (PDB-ID 6ZMW), or an AUG recognition complex with $tRNA_i^{Met}$ in the $P_{IN}$ position/40S head closed (PDB-ID 8PJ1). In all comparisons, structures were aligned on the 18S rRNA of the 40S body. The 18S rRNA, $tRNA_i^{Met}$ and mRNA/circRNA are colored as indicated.

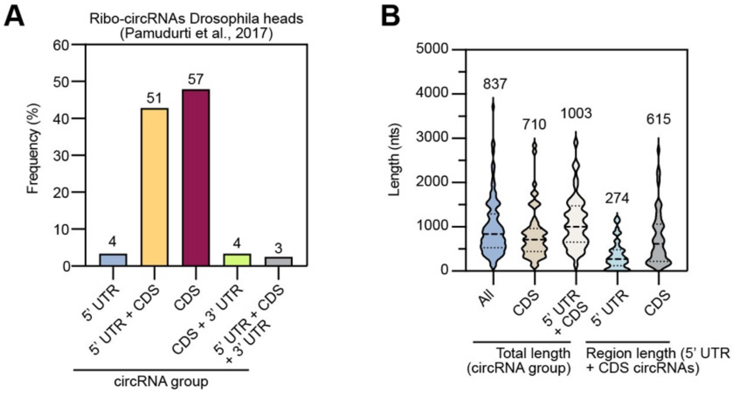


**Figure S6. Features of other ribosome-associated *Drosophila* circRNAs.**

(**A**) Categorization of translated circRNAs from *Drosophila* [22] according to the contained regions (5' UTR, CDS, 3' UTR) in the corresponding exon(s) of their hosting mRNAs. The total number of each category is given above the bars.

(**B**) Total lengths of all, CDS-only or 5' UTR+CDS circRNAs assigned in (A), and length of the 5' UTR or CDS regions with the 5' UTR+CDS circRNAs shown as violin plot. The median lengths is given above each violin.

**Table S1.** Mass spectrometry analysis of circMbl/Sfl-FLAG 48S initiation complexes purified from RRL. A negative control containing no RNA, and a positive control with VHP-β mRNA were used.

**Table S2.** Cryo-EM data collection, refinement, and validation statistics for the circMbl-FLAG scanning and AUG recognition complexes.

| | **circMbl-FLAG scanning complex** (PDB 9SJ4) (EMDB EMD-54937) | **circMbl-FLAG AUG recognition complex** (PDB 9SJ3) (EMDB EMD-54936) |
|---|---|---|
| **Data collection and processing** | | |
| Microscope | FEI Titan Krios G3 | |
| Voltage (keV) | 300 | |
| Camera | Gatan K3 post-GIF direct electron detector + Gatan BioQuantum energy filter | |
| Nominal magnification (x fold) | 105,000 | |
| Pixel size at detector (nominal/calibrated) (Å/pixel) | 0.826 / 0.824 | |
| Physical pixel dose rate ($e^-$/pixel/s) | 18.44 | |
| Exposure time (s) | 1.48 | |
| Total electron exposure ($e^-/Å^2$) | 40 | |
| No. of frames collected during exposure | 111 | |
| No. of fractions grouped | 40 | |
| Dose per fraction ($e^-/Å^2$) | 1 | |
| Defocus range (µm) | -2.8, -2.6, -2.4, -2.2, -2.0, -1.8, -1.6, -1.2 | |
| Automation software | EPU v. 2.12.2 | |
| Micrographs collected (no.) | 16,320 | |
| Processing software | RELION 4.0 and 5.0 | |
| Micrographs used (no.) | 15,388 | |
| Extracted particles total/ribosomal (no.) | 2,272,647 / 156,791 | |
| Final particles (no.) | 21,721 | 79,766 |
| Point-group or helical symmetry parameters | C1 | |
| **Resolution estimates[1] (global, Å) (unmasked/masked)** | | |
| Half-maps FSC = 0.143 | 3.93 / 3.12 | 3.05 / 2.71 |
| Model-map FSC = 0.5 | 4.04 / 3.19 | 3.09 / 2.80 |
| $d_{99}$ | 2.70 / 3.08 | 2.56 / 2.93 |
| $d_{model}$ | 2.80 / 2.90 | 2.80 / 2.80 |
| Resolution range[2] (local, Å) | 2.68 – 22.31 | 2.34 – 18.17 |
| Map sharpening *B* factor ($Å^2$) | 0 | 0 |
| **Model composition** | | |
| Non-hydrogen atoms | 113,724 | 114,857 |
| Protein residues | 10,384 | 10,363 |

| | | |
|---|---|---|
| RNA nucleotides | 1,792 | 1,819 |
| $Mg^{2+}$ ions | 76 | 88 |
| $K^+$ ions | 1 | 3 |
| $Zn^{2+}$ ions | 3 | 3 |
| **Model refinement** | | |
| Refinement package | phenix.real_space_refine_1.21.2-5419 | |
| **Model-Map scores** | | |
| CC (mask) | 0.85 | 0.91 |
| CC (volume) | 0.84 | 0.90 |
| **Average grouped B factors (Å$^2$)** | | |
| Protein residues | 173.22 | 168.16 |
| RNA nucleotides | 89.92 | 89.87 |
| Ligands | 52.32 | 51.83 |
| **r.m.s.d. from ideal values** | | |
| Bond lengths (Å) | 0.005 | 0.008 |
| Bond angles (°) | 0.933 | 1.023 |
| **Validation** | | |
| MolProbity score | 1.69 | 1.61 |
| CaBLAM outliers (%) | 3.75 | 3.20 |
| Clash score | 5.30 | 4.33 |
| Poor rotamers (%) | 0.28 | 0.23 |
| C-beta deviations (%) | 0.02 | 0.01 |
| **Ramachandran plot** | | |
| Favoured (%) | 93.88 | 94.17 |
| Allowed (%) | 5.99 | 5.77 |
| Outliers (%) | 0.13 | 0.06 |
| **Ramachandran plot *Z*-score, (r.m.s.d.)** | | |
| whole | -3.78 (0.07) | -3.56 (0.07) |
| helix | -3.69 (0.05) | -3.54 (0.05) |
| sheet | -0.95 (0.14) | -0.82 (0.13) |
| loop | -2.04 (0.08) | -1.77 (0.08) |

[1] Calculated using Phenix Comprehensive Validation tool (mask based on model)
[2] Calculated using Relion's local resolution estimation
